# Multiplex-GAM: genome-wide identification of chromatin contacts yields insights not captured by Hi-C

**DOI:** 10.1101/2020.07.31.230284

**Authors:** Robert A. Beagrie, Christoph J. Thieme, Carlo Annunziatella, Catherine Baugher, Yingnan Zhang, Markus Schueler, Alexander Kukalev, Rieke Kempfer, Andrea M. Chiariello, Simona Bianco, Yichao Li, Antonio Scialdone, Lonnie R. Welch, Mario Nicodemi, Ana Pombo

**Affiliations:** Epigenetic Regulation and Chromatin Architecture Group, Berlin Institute for Medical Systems Biology, Max-Delbrück Center for Molecular Medicine, Hannoversche Str. 28, 10115 Berlin, Germany; Laboratory of Gene Regulation, Weatherall Institute of Molecular Medicine, Oxford, OX39DU, UK; Dipartimento di Fisica, Università di Napoli Federico II, and INFN Napoli, CNR-SPIN, Complesso Universitario di Monte Sant’Angelo, 80126 Naples, Italy; School of Electrical Engineering and Computer Science, Ohio University, 45701 Athens, OH, USA; Institute of Epigenetics and Stem Cells, Helmholtz Zentrum München – German Research Center for Environmental Health, Munich, 81377, Germany; Institute of Functional Epigenetics, Helmholtz Zentrum München – German Research Center for Environmental Health, Neuherberg, 85764, Germany; Institute of Computational Biology, Helmholtz Zentrum München – German Research Center for Environmental Health, Neuherberg, 85764, Germany; Berlin Institute of Health (BIH), MDC-Berlin, Berlin, Germany; Humboldt-Universität zu Berlin, 10117 Berlin, Germany

## Abstract

Technologies for measuring 3D genome topology are increasingly important for studying mechanisms of gene regulation, for genome assembly and for mapping of genome rearrangements. Hi-C and other ligation-based methods have become routine but have specific biases. Here, we develop multiplex-GAM, a faster and more affordable version of Genome Architecture Mapping (GAM), a ligation-free technique to map chromatin contacts genome-wide. We perform a detailed comparison of contacts obtained by multiplex-GAM and Hi-C using mouse embryonic stem (mES) cells. We find that both methods detect similar topologically associating domains (TADs). However, when examining the strongest contacts detected by either method, we find that only one third of these are shared. The strongest contacts specifically found in GAM often involve “active” regions, including many transcribed genes and super-enhancers, whereas in Hi-C they more often contain “inactive” regions. Our work shows that active genomic regions are involved in extensive complex contacts that currently go under-estimated in genome-wide ligation-based approaches, and highlights the need for orthogonal advances in genome-wide contact mapping technologies.

## Introduction

Our understanding of gene regulation has been dramatically transformed by genome-wide technologies for identifying regulatory elements (e.g. ChIP-seq, ATAC-seq; Andersson and Sandelin, 2020) and by technologies that reveal how these elements are connected to one another through 3D genome conformation (e.g. Hi-C; Kempfer and Pombo, 2020). However, many cell types of interest are too rare to assay using these methods. Although single-cell variants of Hi-C are available, they require purified, disaggregated cell suspensions which can be unachievable for rare cell types embedded within complex tissues. Furthermore, methods based on chromatin conformation capture (3C) usually focus on contacts between pairs of elements, neglecting higher-order associations. We previously showed that Genome Architecture Mapping (GAM) can identify three-way chromatin contacts and achieves strong enrichment for contacts between regions containing active genes, enhancers and super-enhancers whilst requiring only a few hundred cells (Beagrie et al., 2017).

The original GAM protocol involves DNA sequencing of many individual thin nuclear slices (which we call nuclear profiles, or NPs), each isolated in a random orientation from a different cell in the population. The principle behind GAM is that DNA loci which are physically close to each other in the nuclear space are present in the same NP more frequently than loci which are remote from one another. In the prototype version of GAM, a collection of 200-nm cryosections were cut through a sample of mES cells, before microdissection of single NPs into separate PCR tubes, followed by lengthy manual preparation of sequencing libraries to determine the DNA content of each tube.

We have made several significant improvements to GAM. First, to reduce the hands-on time required for sequencing hundreds or thousands of NPs, we developed multiplex-GAM. In this variant of GAM, multiple NPs can be added into a single tube and then sequenced together, cutting down on both labour and reagent costs. Second, we optimized the protocol for DNA extraction from NPs such that it is now compatible with liquid dispensing robots, further reducing time and reagent volumes required to generate a GAM dataset. Third, we extended the SLICE statistical model for analysis of GAM data to cover a wider range of experimental parameters, including the addition of multiple NPs per tube. We also use the SLICE statistical model to determine optimal experimental parameters to aid the design of GAM experiments in different cells and organisms. Fourth, we produce an extended GAM dataset from mES cells, which we use for comparisons with Hi-C. Finally, we show that many contacts are equally detected by both methods, but also identify method-specific contacts, especially those that involve simultaneous associations between three or more genomic elements. We show that GAM is a versatile methodology for mapping chromatin contacts, and provide a framework to design GAM experiments that considers the depth of chromatin contact information required and minimises data collection effort.

## Results

We previously published a GAM dataset of 408 NPs from mES cells, each isolated from a different nucleus into a single PCR tube (Fig. 1a, original-GAM). We showed that the number of times particular genomic loci are found together in the same NP (their co-segregation) is a measure of their physical proximity in the original population of cells, with high co-segregation values indicating that the regions were close in space. In the first GAM dataset, loci on different chromosomes (which we assume will be distant in the nuclear space) are found together in less than 1% of NPs (Beagrie et al., 2017). We therefore reasoned that combining more than one NP into a single sequencing library would not reduce our ability to distinguish interacting from non-interacting loci (Fig. 1a, multiplex-GAM).

**Figure 1.**
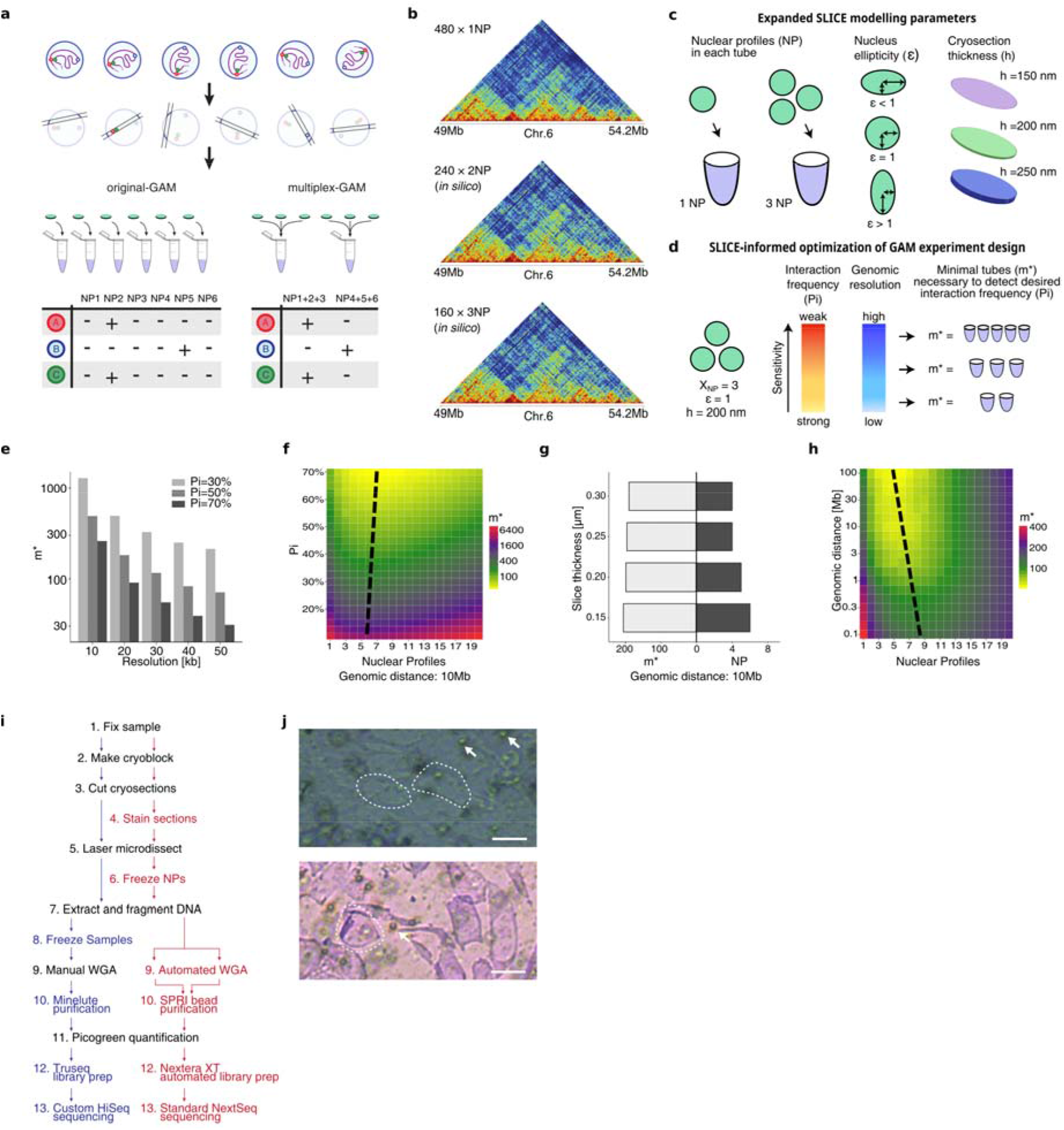
**a**, In a standard GAM experiment, thin slices from individual nuclei (nuclear profiles or NPs) are isolated by cryosectioning and laser microdissection, then the DNA content of each slice is determined by next generation sequencing. In a multiplex-GAM experiment, DNA from multiple NPs is extracted and sequenced together, reducing sequencing costs. **b**, Multiplex-GAM data constructed *in silico* by combining 1NP datasets at 40 kb resolution. **c**, SLICE accounts for the number of NPs multiplexed in each tube (*X_NP_*), the nuclear ellipticity (Supplementary Fig. S1a) and the thickness of each NP (*h*) to estimate the minimal number of samples/tubes (*m\**) needed to achieve a given statistical power. **d**, Relationship between the number of tubes required (*m\**), genomic resolution and sensitivity. **e**, Required number of tubes (*m\**) as a function of genomic resolution and interaction frequency *Pi* given a genomic distance of 10Mb, three NPs per tube (*X_NP_*=3) and detection efficiencies estimated from the previously published GAM dataset at the corresponding resolution (Beagrie et al., 2017). **f**, *m\** as a function of *X_NP_* and the interaction probability (*Pi*) given perfect detection efficiency, a genomic distance of 10Mb and *h*=220nm, at 30 kb resolution. Dashed black line marks the approximate position of the minimum value of *m\** across each row. **g**, Optimal value of *X_NP_* and the corresponding value of *m\** for a range of possible slice thicknesses given *Pi* =30%, genomic distance of 10Mb and perfect detection. **h**, *m*\* as a function of *X_NP_* and the genomic distance given perfect detection, *Pi*=50% and *h*=220nm, at 30 kb resolution. Grey line marks the position of the minimum value of *m\** across each row. **i**, Comparison of the original (black and blue lettering) and multiplex (black and red lettering) GAM protocols. WGA: Whole Genome Amplification. **j**, Top: unstained cryosections from mES cells. Individual NPs are outlined by dashed white lines. White arrows indicate typical background (air bubbles) in the microdissection membrane. Bottom: Cresyl violet stains both cytoplasm and nucleoplasm in mES cell cryosections as visualized by brightfield microscopy, and therefore does not distinguish between cellular profiles that do or do not intersect the nucleus.

To test this intuition, we used a dataset of 481 NPs sequenced individually, which include 408 previously published samples (Beagrie et al., 2017) plus 73 additional NPs (Supplementary Table 1). To simulate multiplex sequencing of two or three NPs, we combined the 481 datasets from single NPs, and generated 240 or 160 *in silico* GAM samples containing two or three NPs, respectively. We then re-calculated co-segregation matrices from these simulated multiplex-GAM datasets and found that these were visually highly similar (Fig. 1b; Supplementary Fig. S1c).

To formally understand the effect of including several NPs in multiplex-GAM experimental designs, and to optimize the experimental parameters, we next updated SLICE, the statistical tool previously developed to infer non-random DNA interaction probabilities from locus co-segregation in GAM data (Beagrie et al. 2017). We considered the effects of number of NPs per GAM sample, nuclear ellipticity, and NP thickness (Fig. 1c, Supplementary Fig. 1a). To determine the optimal parameters for collection of multiplex GAM datasets in mES cells, we expanded the mathematical model SLICE (Beagrie et al., 2017), and estimated the minimal number of GAM samples (*m\**) required to detect chromatin contacts with a probability of interaction above a specific threshold (*Pi*), given the number of NPs per GAM sample (*X_NP_*), nucleus ellipticity, section thickness (*h*) and genomic resolution (Fig. 1d, Supplementary Note). We find that *m\** increases for higher resolutions and for weaker interaction strengths (lower *Pi*), as expected (Fig. 1e,f). We also find that *m\** remains constant over the range of cryosection thicknesses convenient for ultracryomicrotomy, while *X_NP_* decreases with increasing thickness (Fig. 1g). Finally, we show that the expected co-segregation frequency of two interacting loci is relatively invariant for all but the most extremely ellipsoidal nuclei (Supplementary Fig. 1b).

Using the updated SLICE model, we calculate optimal experimental parameters for the application of GAM to a range of different organisms and cell types. Despite differences in ploidy and nuclear geometry between the selected cell types, we find that the minimum number of tubes (*m\**) required to reach a given statistical power is approximately constant (~150 tubes to detect interactions with *Pi* ≥30%) provided that each sample is collected with the optimal number of multiplexed NPs (Supplementary Fig. 1c).

Finally, we determined the optimal experimental parameters for producing a new multiplex-GAM dataset in mES cells, and found that *m\** reaches a minimum with a multiplexing factor of 3-10 NPs per GAM library (Fig. 1h). A GAM dataset of ~250 libraries each multiplexed with 3 NPs (i.e. obtained from a total of only 750 mES cells) would be sufficient to sample contacts with interaction probabilities above 50% at resolution of 30 kb across genomic distances >100 kb, while reducing reagent costs by two thirds (Fig. 1h).

Next, we implemented an improved experimental pipeline for GAM data collection which incorporates staining of the cells for better identification, and shorter or improved steps to facilitate high-throughput experiments, such as storage of laser microdissected NPs, automated pipetting, DNA purification, amplification and sequencing (Fig. 1i). One challenge in the prototype version of GAM was that some cryosections intersect cells without intersecting the nucleus and therefore contain no genomic DNA. In our efforts to identify chemical stains that could allow direct visualisation of nucleus prior to microdissecion, we found that most DNA stains prevent subsequent extraction and/or amplification of DNA from NPs, probably because they bind too strongly or damage DNA (Supplementary Fig. 2a). We therefore collected most of the new multiplex-GAM dataset using a cresyl violet stain which greatly improves identification of cellular profiles during microdissection without affecting DNA extraction (but which does not distinguish the cytoplasm from the nucleus; Fig. 1j, Supplementary Fig. 2b).

In the absence of a direct nuclear stain, we next optimized our multiplex-GAM collection by determining the proportion of cellular slices devoid of NPs. We stained mES cell cryosections for DNA and RNA and found that 23% of profiles intersect the cytoplasm only, with no detectable nuclear material (Supplementary Fig. 2c). As our current GAM pipeline does not distinguish these “cytoplasmic profiles” from NPs, we calculated that if we dissected four cresyl violet stained profiles into each sequencing tube, on average three of those four profiles would intersect the nucleus (Supplementary Fig. 2d).

To directly test the multiplex-GAM approach with our revised experimental pipeline, we collected a new batch of 249 multiplex GAM sequencing libraries, each containing three nuclear profiles on average, from an independent biological replicate from mES cells (Fig. 2a,b). The distribution of quality metrics (most importantly the percentage of mapped reads) was comparable across different collection batches (Supplementary Fig. 3a). The genomic coverage of this dataset (18% of 40 kb windows are detected per NP on average) was consistent with the expected presence of three NPs per library on average (7% for single NPs, 20% for 3NP *in silico* data; Supplementary Fig. 3b) (Beagrie et al., 2017). Comparison of normalized linkage matrices between the 249×3NP multiplex-GAM dataset and the 481×1NP original-GAM dataset indicated that local contact information is well preserved in multiplex-GAM, and conserved between biological replicates (Fig. 2a,b). Erosion analysis of the 3NP-GAM dataset at 40 kb shows that its correlation with the 1NP-GAM dataset resolution begins to saturate at 200×3NPs, indicating that the 249×3NP dataset is large enough to explore chromatin structure at this resolution (Supplementary Fig. 3c).

**Figure 2.**
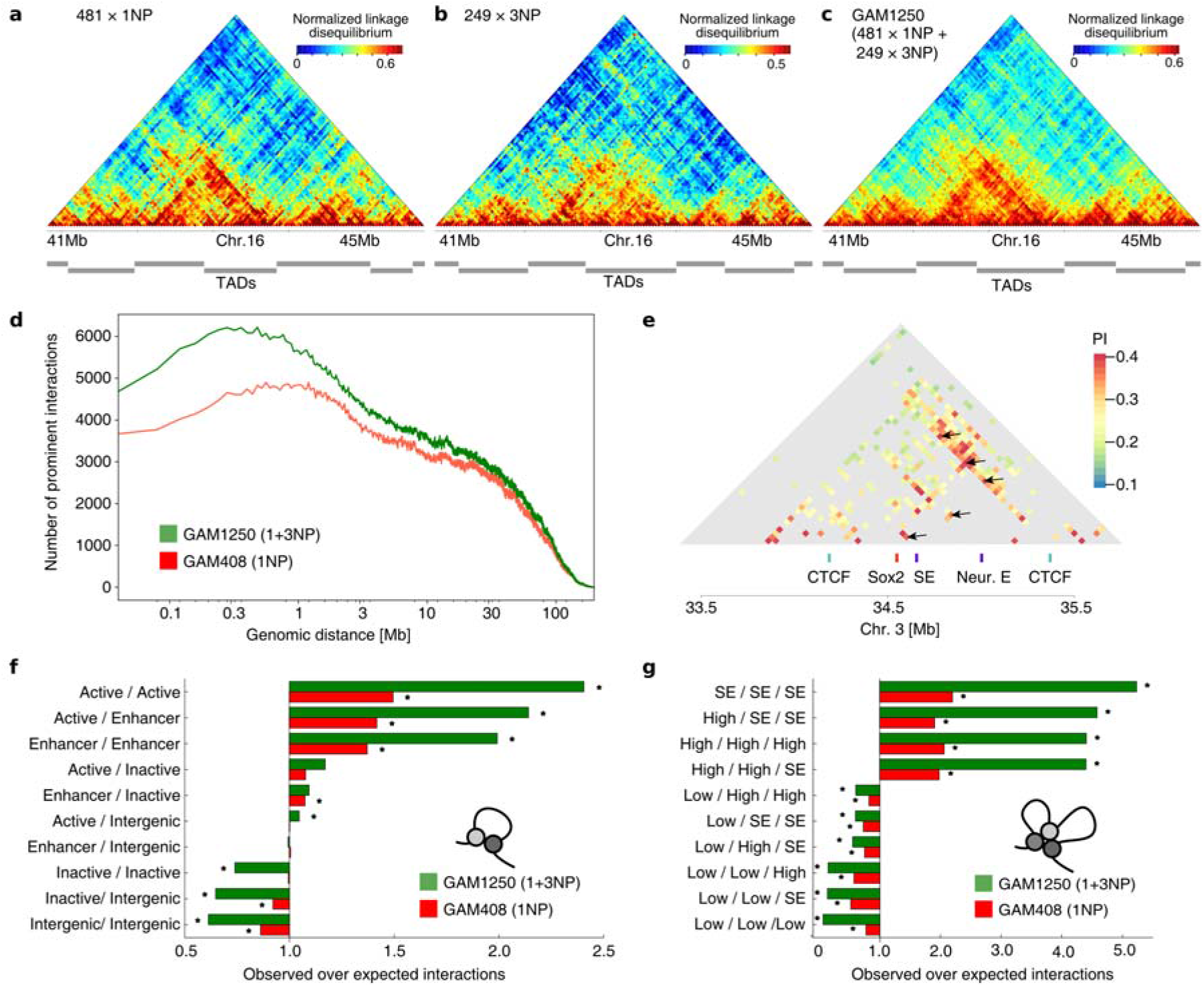
GAM matrices of normalized linkage with a resolution of 40 kb for **a**, multiplex-GAM using 3NPs (249 datasets), **b**, GAM using 1NPs (481 datasets), and **c**, the combined dataset of 1NPs and 3NPs. **d**, Number of prominent interactions identified by SLICE at each genomic distance in the merged GAM-1250 dataset and the published mES GAM dataset (Beagrie et al., 2017). **e**, Contact matrix centered at the Sox2 locus presenting prominent interactions identified by SLICE at 40 kb resolution. Functional elements in the locus are identified below the matrix, and previously identified interactions between these elements are identified by black arrows (Bonev et al., 2017). **f**, Enrichment analysis of pairwise interactions identified by SLICE involving active, inactive, intergenic or enhancer regions for the merged GAM-1250 dataset (green) and the original GAM-408 dataset (red). Statistically significant enrichments/depletions (those falling outside 95% of randomised observations after Bonferroni correction) are marked by an asterisk. **g**, Enrichment analysis of triplet interactions identified by SLICE for the merged GAM-1250 dataset (green) and the original GAM-408 dataset (red).

We next considered the possibility of merging the 1NP and 3NP datasets to increase our power for in-depth comparisons between GAM and Hi-C data. First, we used the 1NP-GAM data to test *in silico* the effect of combining 1NP and 3NP data, and found that matrices from combined 1NP and 3NP libraries retain the quality of 3NP libraries (Supplementary Fig. 4a). Therefore, we merged the experimental 1NP and 3NP datasets to create a combined GAM dataset spanning a total of 1250 NPs, each from a different cell (Fig. 2c). To confirm the increased statistical power of the combined 1+3NP dataset, we used SLICE to determine the prominent interactions and compared them with those obtained with the original 408×1NP dataset (Beagrie et al., 2017). Using the deeper 1250 NP dataset from combined 1+3NP GAM data, we detect a greater number of prominent interactions compared to the published data (Fig. 2d,e). These prominent interactions showed a similar pattern of enrichment for genomic features as previously shown in our original 408×1NP mES cell dataset containing only 1NP samples, both for pairwise (Fig. 2f; Supplementary Fig. 4b) and three-way interactions (Fig. 2g).

One of the key aims of genome-wide 3D chromatin folding assays is the detection of TADs (Beagan and Phillips-Cremins, 2020). We used the insulation score method (Crane et al., 2015) to identify TAD boundaries in GAM and published Hi-C data (Dixon et al., 2012), choosing parameters that maximized overlap with published Hi-C TAD boundaries (Methods; Fig. 3a; Supplementary Fig. 5a-d). Of the boundaries identified by GAM, 95% matched a boundary identified by Hi-C, whilst 76% of Hi-C boundaries matched a boundary in GAM (Fig. 3b), comparable to the overlap of 60% for boundaries detected from the same Hi-C data using two different computational algorithms (Fig. 3c).

**Figure 3.**
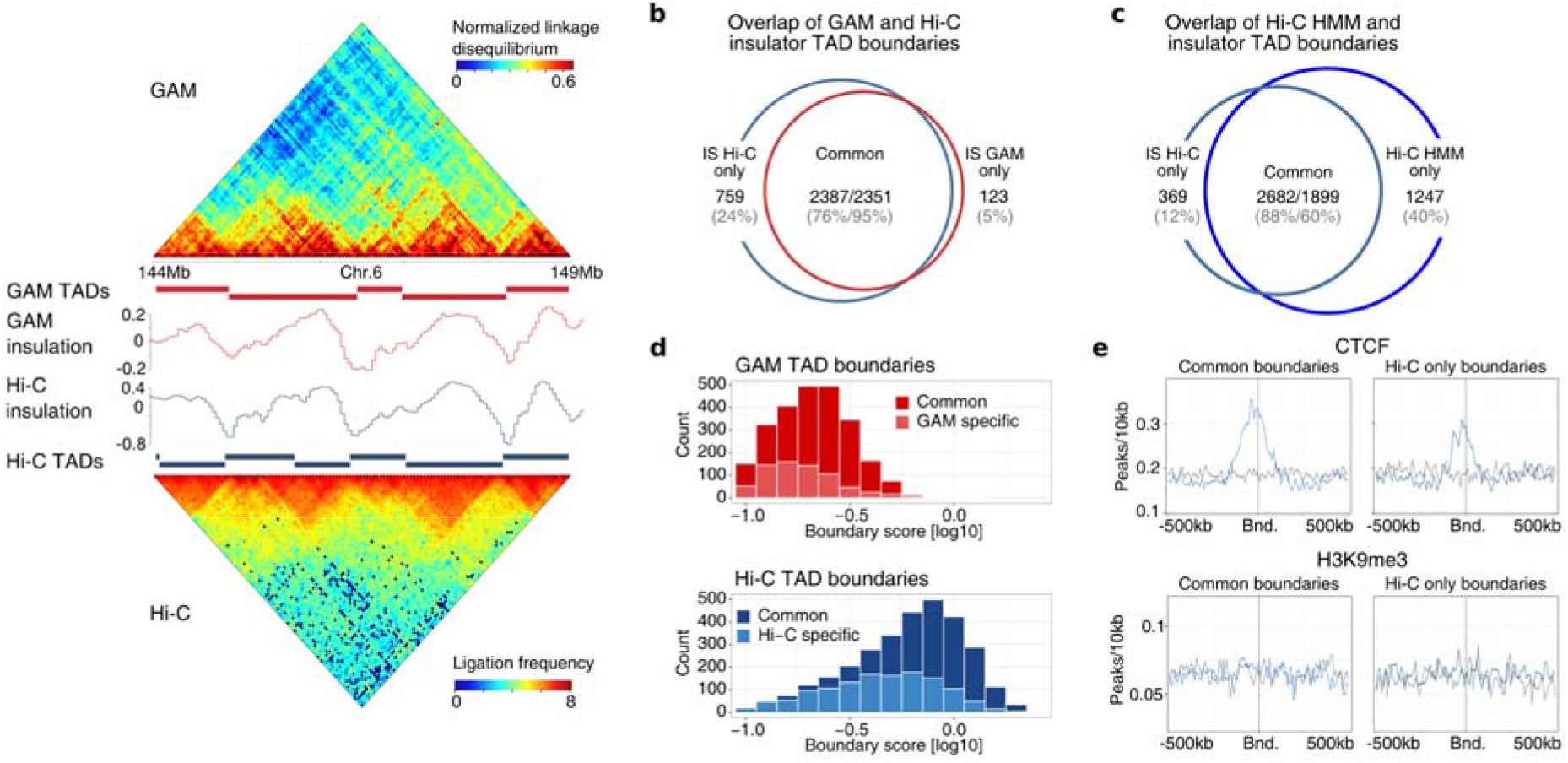
**a**, Comparison Topological associated domains (TADs) defined by the insulation score method in GAM (top) and Hi-C (Dixon et al., 2012; bottom). **b**, Overlap (within 80 kb distance) of TAD boundaries detected in GAM (red) and Hi-C (blue). **c**, Overlap between insulator score and Hidden Markov Model (HMM) TAD boundary calls from the same Hi-C data. **d**, Boundary strength (drop of insulation score) measured by GAM (red) and Hi-C (blue) for TAD boundaries that are shared between the two methods (dark bars) or only found in a single method (light bars). **e**, Enrichment (blue) of CTCF and H3K9me3 around common or Hi-C-specific TAD boundaries (Bnd.), compared to genomic background using shuffled boundary positions (grey). Enrichments for GAM-only boundaries are not shown as these are very few.

We compared the strength of boundaries detected by only one method with those detected by both GAM and Hi-C. We find that TAD boundaries identified by Hi-C but not GAM (750 boundaries), or by GAM but not Hi-C (123 boundaries), have weaker insulation than common boundaries (Fig. 3d). To investigate whether the boundaries that are differentially detected between Hi-C and GAM have any distinctive features, we plotted the enrichment of various chromatin marks over TAD boundaries. We confirm the enrichment of previously described features at common boundaries (e.g. common boundaries are enriched for CTCF binding, H3K36me3 and SINE elements, but not for H3K9me3; Fig. 3e; Supplementary Fig. 5e). In most cases, features found at common boundaries are also found at boundaries identified only in GAM or only in Hi-C except with lower enrichment, again suggesting that the strongest boundaries are detected by both methods. The exceptions to this are H3K4me3, RefSeq TSSes, unphosphorylated RNA Polymerase II and CTCF, for which there was a small enrichment at Hi-C only boundaries but not at the smaller number of GAM-only boundaries (Supplementary Fig. 5e). TAD boundaries identified by Hi-C but not by GAM may therefore be a consequence of the fact that GAM detects more contacts between TADs and at longer distances, consistent with the lower Hi-C insulation score of these boundaries and the lack of any specific molecular features distinguishing them from common boundaries.

To investigate genome-wide differences between GAM and Hi-C in an unbiased fashion, we developed a method for directly comparing matrices derived from the two methods. For these analyses, we considered contacts between loci separated by ≤ 4Mb, as the fidelity of both datasets decreases at larger genomic distances, and our simulated multiplex-GAM dataset confirmed that most of the contact information in pairwise matrices is well preserved within this distance when combining NPs in sets of three (Supplementary Fig. 1d). The selected genomic length scale is useful in most current applications of chromatin contact mapping, in particular, it is sufficient for the detection of enhancer-promoter contacts in most instances.

Because GAM detects far more contacts at larger genomic distances (Kempfer and Pombo, 2020) and because data generated by the two techniques has very different numerical distributions, it remained challenging to identify regions with similar or divergent contact patterns between GAM and Hi-C. GAM normalized linkage values of genomic window co-segregation range from −1 to +1, whereas Hi-C ligation frequency is dependent on sequencing depth and ranges over many orders of magnitude. We therefore first applied a distance-based z-score normalisation to both datasets to address the distance-decay (Fig. 4a). Next, we subtracted the two normalized matrices and extracted the most divergent contacts where the difference between the two matrices was greater than the 5% extremes defined by a fitted Normal distribution. We refer to these most divergent contacts as GAM-specific or Hi-C-specific contacts (Fig. 4a).

**Figure 4.**
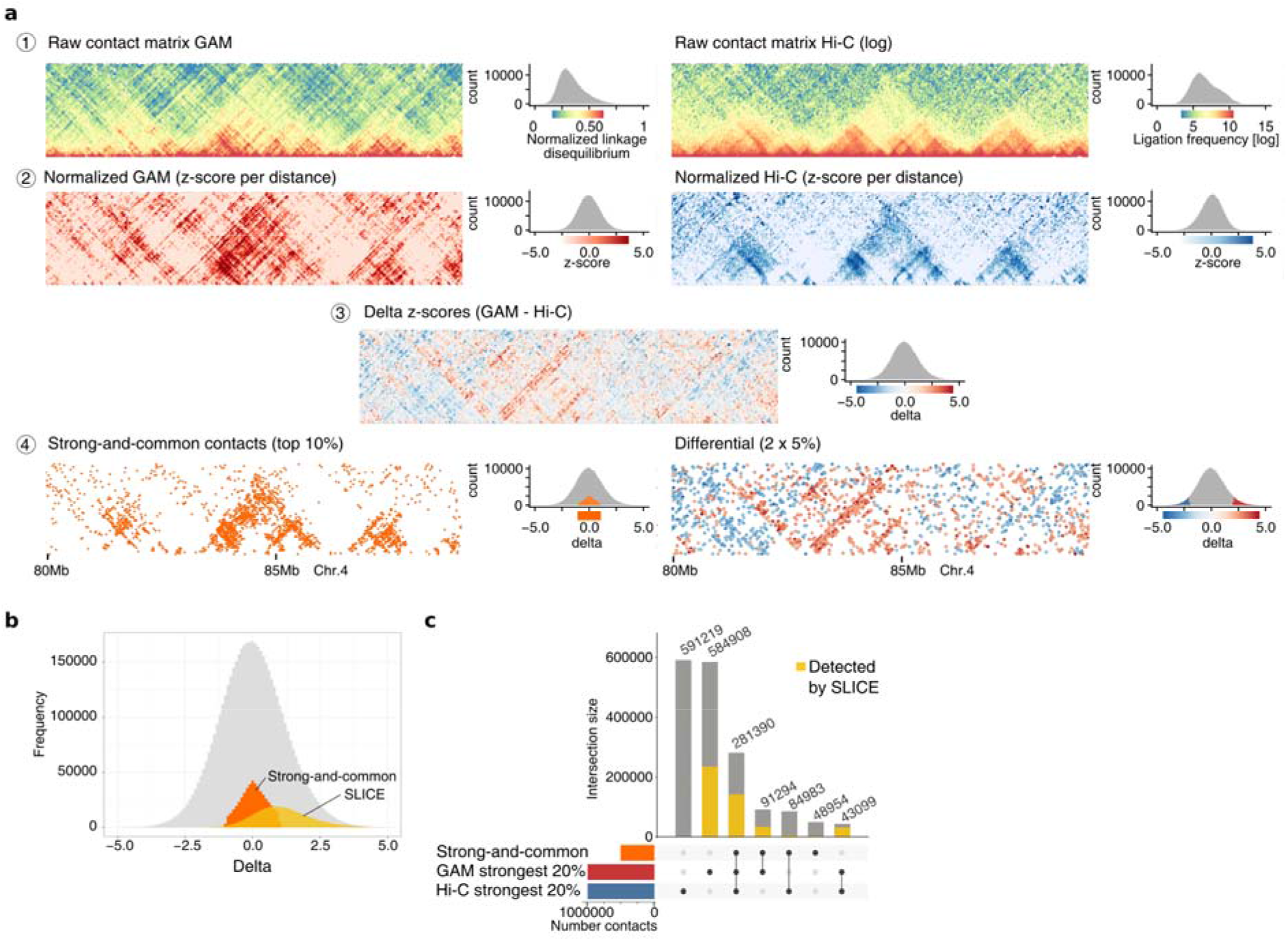
**a**, Strategy to assess differences and similarities between GAM and Hi-C contact maps. GAM and Hi-C contact data have very different distributions (1), so contacts at the same genomic distance are normalized by z-score transformation (2). Hi-C z-scores are then subtracted from GAM z-scores to generate a delta z-score matrix (3), from which we extract the top 10% of contacts common for both methods (strong-and-common; 4) and the most differential contacts between GAM and Hi-C (GAM-specific or Hi-C specific; 5). **b**, Prominent interactions identified by SLICE shown within the distribution of delta z-scores. **c**, Set sizes for contacts found in combinations of the top 20% strongest contacts from GAM and Hi-C, and the strong-and-common set. Split bars indicate fraction of each set supported by SLICE prominent interactions.

Next, we established a set of strong-and-common contacts that are well detected in both methods by taking the 10% strongest contacts which showed a z-score delta of less than 1.0. Many of the strong-and-common contacts are also prominent interactions detected by SLICE analyses of GAM data (Fig. 4b). Strong-and-common contacts and SLICE interactions were more frequently separated by genomic distances below 2 Mb (Supplementary Fig. 6a), whereas GAM-specific and Hi-C-specific contacts were slightly depleted at distances <1Mb (Supplementary Fig. 6b). To verify that the GAM-specific and Hi-C-specific contacts are indeed strong contacts in each corresponding method (i.e. they are not artefacts of our normalisation procedure), we compared their raw intensities. These analyses confirmed that our approach successfully identifies contacts with high intensity in GAM but low intensity in Hi-C (or vice-versa; Supplementary Fig. 6c). Importantly, whilst many of the strongest contacts in absolute terms were identified by both methods, more than half of the strongest 20% of contacts in GAM or Hi-C were method specific (Fig. 4c).

To investigate whether the contacts differentially detected by GAM or Hi-C hold important biological roles, we asked whether they were enriched for particular genomic features. We created a dataset of features including repeat elements, heterochromatin marks, transcription factor binding sites, RNA Polymerase II and transcription-related histone marks (Fig. 5a). We then counted the number of contacts in each category (GAM-specific, Hi-C-specific, strong-and-common) between each possible pair of features (e.g. CTCF-CTCF, EP300-Nanog, etc.), and looked for feature pairs over-represented (enriched) or under-represented (depleted) from GAM-specific or Hi-C-specific contacts relative to distance-matched random backgrounds (Methods; Supplementary Fig. 7a). We found many feature pairs enriched in GAM-specific contacts, whereas only a small number were enriched in Hi-C-specific contacts (Supplementary Fig. 7b,c). The feature pairs most enriched in GAM-specific contacts included transcription factors, enhancers and RNA Polymerase II (Fig. 5b). By contrast, heterochromatin regions (i.e. those marked by H3K9me3 or H3K20me3) were most enriched in Hi-C-specific contacts.

**Figure 5.**
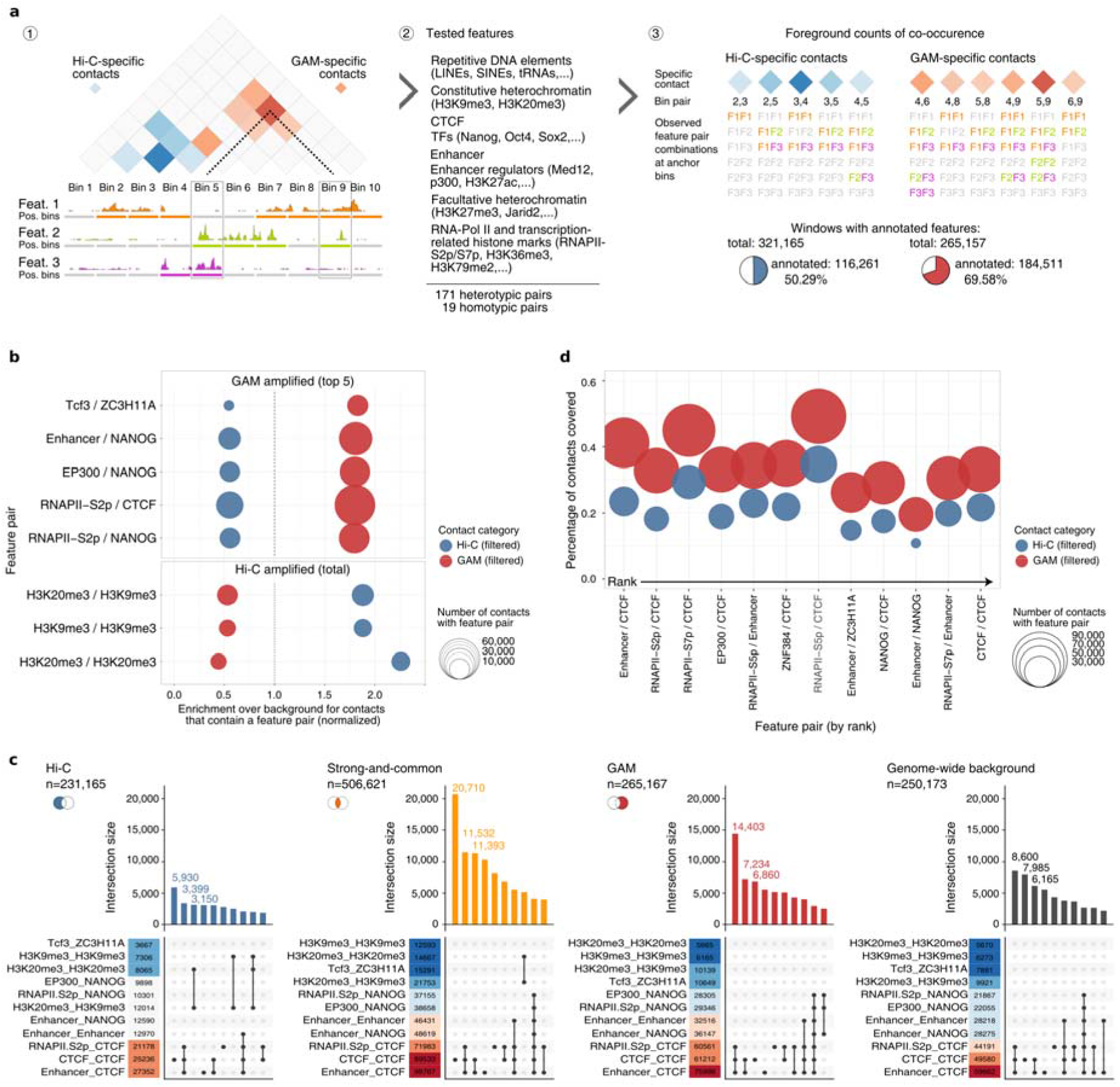
**a**, Schematic for detecting feature pair enrichments in method-specific contacts (red and blue). (1) Each contact is defined by two genomic anchor points which we categorized as either positive or negative for peaks of the respective feature. (2) We assessed 190 feature pairs and (3) quantified feature occurrences at the anchor points, filtering out contacts with no feature pairs. **b**, Feature pairs with the highest enrichment over background for GAM-specific (top) and Hi-C-specific (bottom) contacts. **c**, Upset plots quantifying the co-occurrences of enriched feature pairs in contacts of each subgroup. Occurrence counts for each feature pair are coloured according to their absolute frequency within contacts of each set. **d**, Feature pairs with the highest discriminatory power (by mean decrease of Gini impurity) in random forests trained to separate GAM-specific and Hi-C-specific contacts.

As this analysis is conducted at 40 kb resolution, many contact anchors may contain multiple genomic features. We therefore stratified our analysis to account for overlapping feature pair annotations (Fig. 5c). For example, the GAM-specific, Hi-C-specific and strong-and-common contact sets all contain large numbers of CTCF-CTCF contacts, being enriched over the genome-wide background in the strong-and-common set and GAM-specific contacts, but depleted in Hi-C-specific contacts. Amongst the GAM-specific contacts, the most common subgroup of CTCF-CTCF contacts consisted of those in which contact anchors also overlapped enhancers and the elongating form of RNAPII (a marker of active genes; 14,403 of 61,212 total CTCF-CTCF contacts). By contrast, in Hi-C-specific and strong-and-common contacts, the most common CTCF-CTCF contacts were those without any other annotation (Hi-C specific: 5,930 of 25,236 total CTCF-CTCF contacts; strong-and-common: 20,710 of 89,533). These results suggest that GAM-specific contacts are strongly enriched for a specific subset of CTCF-CTCF contacts that co-occur with enhancer and active genes and which are underestimated in Hi-C data. Indeed, the enhancer-CTCF class of contacts was the most informative in a random-forest model trained to distinguish GAM-specific and Hi-C specific contacts (Fig. 5d).

Having identified striking enrichments for specific genomic features amongst GAM- and Hi-C-specific contacts, we wondered whether specific features might be generally poorly detected in contacts specific to either method. To identify blind-spots in each method, we developed an approach that counted the number of GAM-specific and Hi-C-specific contacts formed by each window and looked for regions involved in many more GAM-specific than Hi-C-specific contacts or vice versa (Fig. 6a; Supplementary Fig. 8a). Surprisingly, we find that blind-spot windows are not only abundant but also cluster in the linear genome, such that large regions consistently interact more strongly either in GAM or in Hi-C (Fig. 6b). Clusters that form many GAM-specific contacts (GAM-preferred regions) contained more genes and had higher transcriptional activity (Fig. 6c) whilst clusters that form many Hi-C-specific contacts (Hi-C preferred regions) were more frequently found in regions associated with the nuclear lamina (Fig. 6d). We further identified that GAM-preferred regions tend to be occupied by CTCF, P300, certain mES-cell TFs, RNA Polymerase II (especially the elongating, S2p form), enhancers and super-enhancers, whereas Hi-C-preferred regions showed a slight enrichment for the heterochromatin marks H3K20me3 or H3K9me3 (Fig. 6e; Supplementary Fig. 8b).

**Figure 6.**
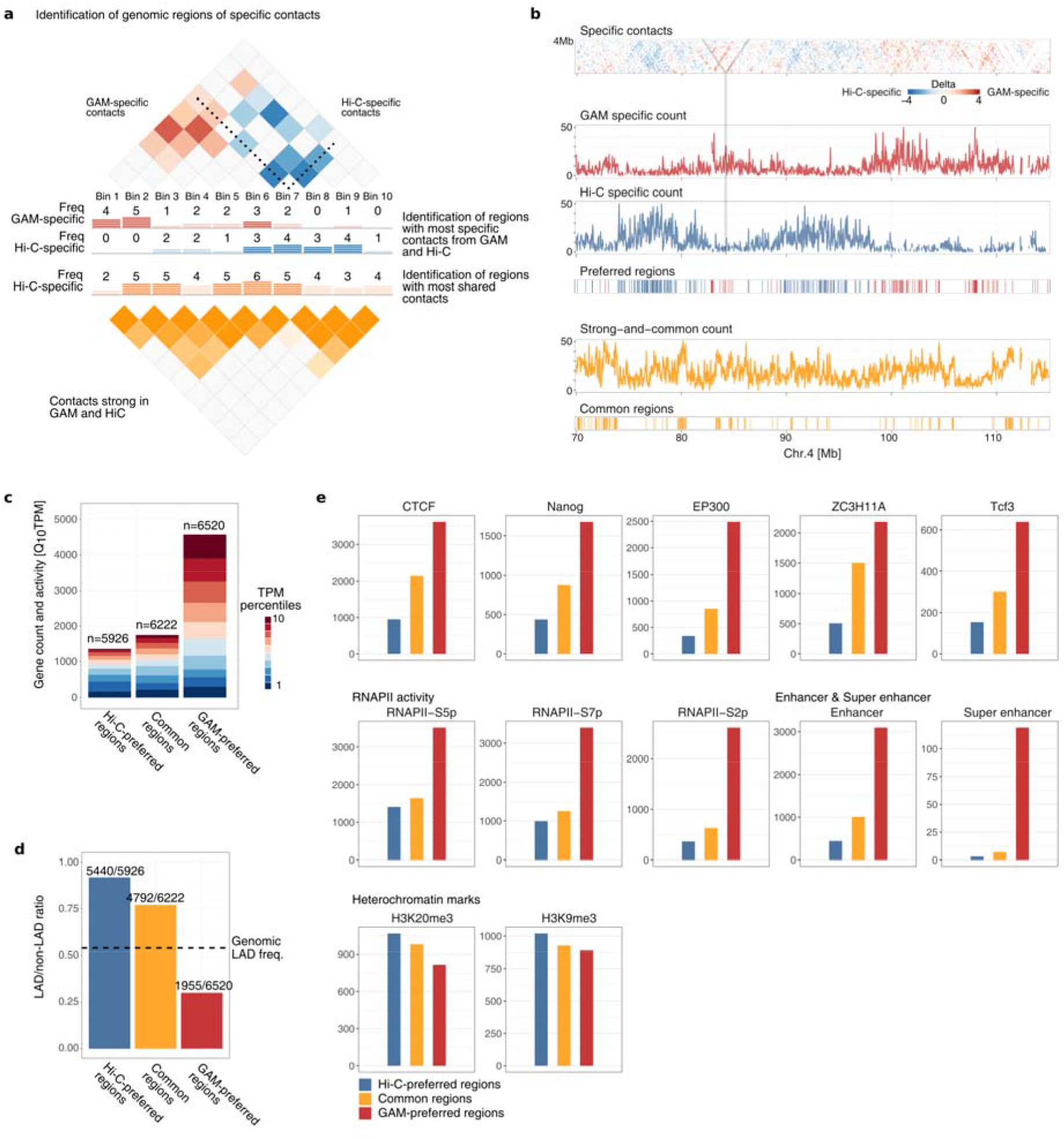
**a**, Strategy for identifying genomic regions forming many contacts specific to either GAM or Hi-C. We counted how often a genomic region was found to be an anchor point in the set of GAM-specific contacts, Hi-C-specific contacts, or strong-and-common contacts. The 10% of genomic windows with the highest absolute difference between the number of GAM-specific and Hi-C-specific contacts were classified as Hi-C-preferred regions or GAM-preferred regions, respectively. Similarly, top 10% of strong-and-common contacts were used to define common regions seen equally well by both methods. **b**, Example region on chromosome 4 showing delta z-score matrix (top) and clusters of method-specific preferences designated as Hi-C-preferred regions, GAM-preferred regions or common regions (bottom). **c**, Number of genes and distribution of gene transcriptional activity (TPM) for Hi-C-preferred (left), common (middle) and GAM-preferred (right) regions. **d**, Fraction of genomic windows within Lamina Associated Domains (LADs) for Hi-C-preferred (left), common (middle) and GAM-preferred (right) regions. Numbers on top of the bar give absolute numbers for LAD windows over the total of windows per category. **e**, Numbers of genomic windows overlapping a range of features.

Since the observed patterns can be partly explained by compartmentalisation, we wondered why GAM and Hi-C might measure different relative contact strengths for these compartments. First, we computed compartment definitions from GAM and Hi-C data. Next, we measured the proportion of GAM- or Hi-C-preferred regions that belong to compartments A or B (the active or inactive compartments, respectively), and found that GAM-preferred regions are highly enriched for chromatin belonging to compartment A, whereas Hi-C-preferred regions and strong-and-common regions are enriched within the B compartment (Fig. 7a; Supplementary Fig. 9).

**Figure 7.**
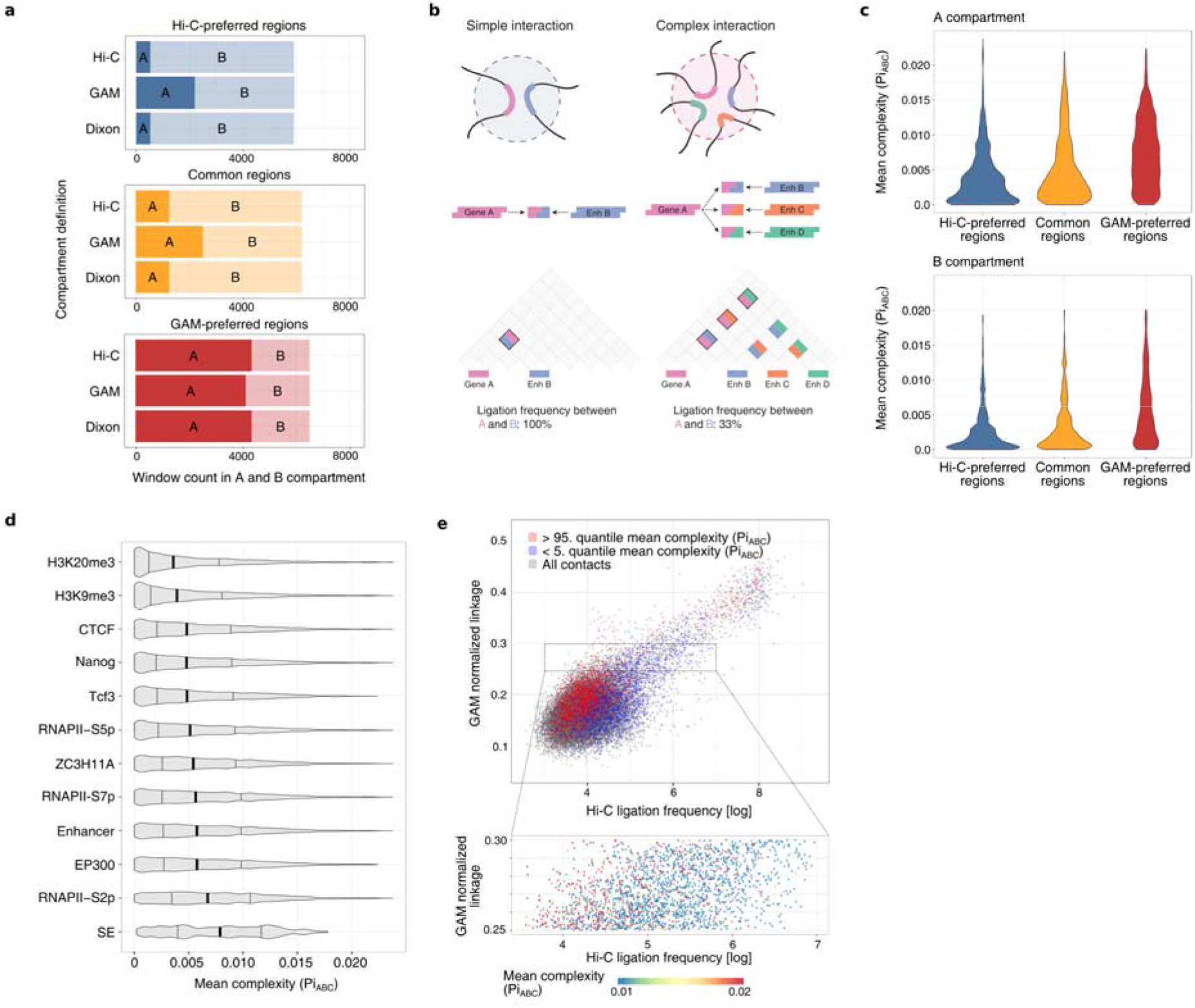
**a**, Number of windows designated as A or B compartments by PCA decomposition of Hi-C data or GAM data, or from published Hi-C PCA compartments (Dixon et al., 2012) in Hi-C-preferred (top), common (middle) and GAM-preferred (bottom) regions. **b**, Interaction complexity: simple interactions involve only a few genomic regions, whilst complex interactions involve many genomic regions at once. Ligation events connect two pieces of DNA, so pairwise contacts can be directly determined while measurement of higher order contacts may be affected by competing ligation events. **c**, Contact complexity (measured as the mean Pi_ABC_ – the interaction probability of 1Mb triplets calculated by SLICE) for Hi-C-preferred (left), common (middle) or GAM-preferred (right) regions in the A compartment (top) or the B compartment (bottom). **d**, Complexity for genomic windows which are positive for a feature, ranked by increasing mean. Vertical bars mark the 25%, 50% and 75% quantiles. **e**, Relationship between pairwise Hi-C ligation frequency (X-axis), pairwise GAM normalized linkage (Y-axis) and complexity (5% pairwise interactions with highest mean *Pi_ABC_* in red, lowest 5% in blue). Inset: Hi-C ligation frequency of 1Mb pairwise contacts that have a similar GAM normalized linkage, coloured by complexity.

Finally, we considered whether the enrichment in GAM-preferred regions for active features (active genes, TFs, polymerase, enhancers and compartment A), could be due to different levels of contact complexity, i.e. to interactions with many simultaneous interacting partners (Fig. 7b). Complex interactions are expected to be underestimated in Hi-C datasets because the ligation step only allows for the measurement of two interacting partners per restriction fragment in each cell where the contact is established (O’Sullivan et al., 2013).

To investigate the relationship between interaction complexity and the differences in method-specific contacts and method-preferred regions, we used SLICE to predict the probability of interaction for all possible sets of three 1Mb windows lying on the same chromosome (i.e. the *Pi_ABC_* for all possible triplets of loci A, B and C; Beagrie et al., 2017). We then calculated the mean *Pi_ABC_* across all triplets involving a particular window. Mean *Pi_ABC_* measures the number of distinct interactions formed by each window that have at least two other partners, and is therefore a measure of interaction complexity. We find that windows in the A compartment indeed form more complex interactions than windows in the B compartment (Fig. 7c). Interestingly, windows that formed many GAM-specific contacts had a much higher complexity than windows that establish strong-and-common or Hi-C-specific contacts even when comparing within the same compartment. All the genomic features associated with GAM-specific contacts were also associated with greater interaction complexity, with the strongest effect seen for the elongating, S2-phosphoisoform of RNA Polymerase II and for super-enhancers (Fig. 7d), in line with our previous work which identified long-range chromatin contacts between super-enhancers and actively-transcribed genomic regions across tens of megabases (Beagrie et al., 2017). These results suggest the existence of abundant chromatin contacts where many active regions interact simultaneously, which have been so far unappreciated through the use of ligation-based methods, but detected by GAM and FISH (Beagrie et al., 2017).

Finally, we examined whether high interaction complexity might artificially deflate pair-wise contact probability as measured by Hi-C, since each DNA fragment picks up a given interacting fragment with lower probability in complex compared with simple contacts (Fig. 7b). We correlated pairwise contacts from GAM and Hi-C at 1Mb resolution and found that regions with a high complexity generally have higher pair-wise contact scores when measured by GAM, whereas regions of low complexity generally have higher pair-wise contact scores when measured by Hi-C (Fig. 7e). This is particularly evident when plotting only a subset of interacting pairs that all have very similar GAM contact frequency (Fig. 7e, zoomed inset), demonstrating that complexity explains a large amount of the divergence in contact frequencies measured by Hi-C or GAM.

## Discussion

The three-dimensional structure of the nucleus is inextricably linked with its functional roles, including gene regulation, DNA replication and the DNA damage response. Consequently, molecular techniques for measuring the 3D folding of chromatin within the nucleus have been instrumental in advancing our understanding of nuclear function over the past decade (Kempfer and Pombo, 2020). Here, we have developed multiplex-GAM, a new variant of Genome Architecture Mapping that enables faster and more cost-effective analysis of chromatin folding genome-wide than the previous version (Beagrie et al., 2017). GAM requires only a few hundred cells, which is of particular relevance to human genetics where researchers need to assay the 3D contacts made by disease-linked sequence variants in specific, often rare cell types impacted by the disease (e.g. neuronal subtypes in neurodegenerative diseases). We have recently applied Multiplex-GAM to different neuronal subtypes in brain tissues, and discovered abundant cell-type specific contacts that contain differentially expressed genes and accessible regulatory elements (Winick-Ng et al., 2020).

Here, we develop multiplex-GAM to generate a larger GAM dataset, and expand the mathematical model SLICE by incorporating new experimental parameters (number of NPs per sample and nuclear ellipticity). We use the larger GAM dataset containing information from 1250 cells for detailed comparisons of the contacts captured by GAM and Hi-C, the most frequently used genome-wide method for chromatin contact analysis (Lieberman-Aiden et al., 2009). We find that GAM and Hi-C detect similar TADs, large folded domains that are thought to constrain gene regulatory elements and form a fundamental unit of chromatin organisation (Dixon et al., 2012; Nora et al., 2012; Sexton et al., 2012). Many strong contacts, including a large proportion of CTCF-mediated loops, are also detected by both methods. By careful examination of finer-scale differences, we identify that chromatin contacts given more weight by GAM frequently connect genomic loci bound by enhancers, key mES-cell transcription factors, RNA Polymerase II and CTCF, whereas contacts which feature more prominently in Hi-C matrices connect regions marked by the heterochromatin-associated histone modifications H3K9me3 and H3K20me3.

We then looked for regions of the genome which consistently form more contacts in GAM datasets than in Hi-C datasets and found that these regions are located in large genomic regions bound by the same activating transcription factors identified in the GAM-specific contacts. In our previous work, we found that super-enhancers are the genomic regions most enriched in complex, multi-partner interactions, together with the most actively transcribed regions (Beagrie et al., 2017). We now extend this finding to show that the regions forming stronger contacts in GAM than in Hi-C tend to form more complex interactions. Theoretical work has previously suggested that ligation-based methods, such as Hi-C, might underestimate contacts between multiple partners, as ligation only captures two or a few contact partners at a time (O’Sullivan et al., 2013). We find that ligation frequencies measured by Hi-C are systematically lower between regions that form complex interactions, providing the first experimental evidence to support such an effect.

Ligation is not the only potential source of difference between the two methods, as GAM and Hi-C also make use of quite different fixation protocols. The choice of fixation protocol has been shown to affect the proportion of informative ligation events between different 3C experiments (Oudelaar et al., 2017), and it may also influence the contacts of genomic regions with different protein composition and/or compaction within a single experiment (Downes et al., 2021; Williamson et al., 2014). The digestion of nuclear chromatin necessary for preparing Hi-C libraries has also been shown to disrupt nuclear structure (Gavrilov et al., 2013), whereas GAM uses fixation protocols specifically chosen to maximize the preservation of nuclear architecture and retention of nuclear proteins (Guillot et al., 2004). Ultimately, formaldehyde fixation remains a “black box” and will continue to complicate interpretation of the most widely used methods for measuring chromatin structure (including microscopy methods like FISH; Gavrilov et al., 2015). Live-cell imaging methods circumvent the need for fixation and will provide valuable orthogonal data, but these methods currently require recruitment of large numbers of fluorophores which may themselves influence folding (Shaban and Seeber, 2020). Ultimately, it should eventually be possible to shed light on the effect of fixation by extending GAM to unfixed nuclei through sectioning of vitrified samples.

Another factor that may influence method-specific contacts is data processing. It has recently been shown that Hi-C detects fewer contacts between regions of condensed chromatin due to a lower accessibility of these regions to restriction enzyme digestion (Chandradoss et al., 2020). However, matrix-balancing algorithms commonly used to normalize Hi-C data can overcorrect for this effect, leading to an aberrantly high frequency of contacts between condensed domains. Consistent with these results, we find that regions of the genome which consistently form more contacts in normalized Hi-C are enriched for heterochromatin marks. We have found the bias in raw GAM datasets to be uniformly lower than that found in raw Hi-C, yet both methods have their own specific biases and improved normalisation algorithms have the potential to bridge the divergences between the two methods (Beagrie et al., 2017; Chandradoss et al., 2020; Kruse et al., 2020; Liu and Wang, 2019).

Our work underscores previous findings that complex, simultaneous interactions between many partners are a pervasive and under-studied feature of mammalian genome architecture (Beagrie et al., 2017; Quinodoz et al., 2018), although their overall prevalence is still a subject of debate (Olivares-Chauvet et al., 2016). Enhancer-binding transcription factors and RNA Polymerase II have both been reported to form nuclear clusters that could serve as nucleating agents for such multi-partner interactions (Iborra et al., 1996; Liu et al., 2014). More recently, there has been a surge of interest in phase-separated nuclear bodies, which are suggested to facilitate high local concentrations of chromatin-interacting proteins and/or transcriptional regulators (Yoshizawa et al., 2020). The clear expectation is that these condensates should bring together multiple interacting genomic partners, in much the same way as rDNA repeats are brought together in the nucleolus (Pederson, 2011). Heterochromatin has also been reported to form phase-separated condensates (Larson et al., 2017), yet we find these regions to have lower-complexity specific interactions, potentially highlighting a shorter-range role for these interactions.

In conclusion, our development of multiplex-GAM, an improved protocol for rapid, cost-effective generation of GAM datasets enabled us to obtain a deeper GAM dataset for mES cells and to explore the similarities and differences between GAM and Hi-C in greater detail. Reassuringly, the two methods paint broadly similar pictures of nuclear architecture, in particular the distribution of TADs, the segregation of nuclear chromatin into A and B compartments and the importance of CTCF for shaping chromatin interactions. There are differences however, with GAM detecting more and stronger contacts between active chromatin regions, and across longer distances, and Hi-C emphasising contacts within silent chromatin. These results highlight the utility of GAM for studying contacts of potential gene-regulatory functions, particularly in human disease where such contacts may only be formed in rare cell populations inaccessible to population Hi-C.

## Methods

### Updated GAM protocol

mES cells were grown and cryoblocks prepared as previously described (Beagrie et al., 2017). 220 nm (green) cryosections were cut with glass knives using a Leica FC7 ultracut cryotome, collected in sucrose droplets (2.1 M in PBS) and transferred to steel frame PEN membrane slides (Leica) UV treated for 45 min prior to use. Slides were washed in sterile-filtered (0.2 μm syringe filter) 1x PBS (3 times, 5 min each), then with sterile-filtered water (3 times, 5 min each). Cresyl violet staining was performed with sterile-filtered cresyl violet (1 % w/v in water, Sigma-Aldrich, C5042) for 10 min, followed by 2 washes with water (30 s each) and air dryed for 15 min. Nuclear profiles were laser microdissected into adhesive 8-strip LCM collection caps (Zeiss), with four profiles dissected into each cap. Caps were stored at −20 °C until ready to proceed with whole genome amplification.

Whole genome amplification of DNA from microdissected NPs was performed using the Sigma WGA4 kit using a liquid handling robot (Microlab STARlet, Hamilton). 14.5 μl of lysis and fragmentation mastermix (13 μl H_2_O, 1.4 μl lysis and fragmentation buffer, 0.09 μl proteinase K) was added to each well of a 96 well plate, caps with microdissected material were used to close the wells, then the plate was inverted and centrifuged upside-down at 3000 xg for 2 min such that the fragmentation mastermix was collected in the cap. Plates were incubated upside-down for 4 h at 50 °C then inverted and centrifuged right-way-up at 3000 xg for 2 min to collect the extracted DNA in the bottom of the well. Samples were then heat inactivated at 99 °C for 4 min then cooled on ice for 2 min. 4.95 μl library preparation mastermix (3.3 μl library preparation buffer, 1.65 μl library stabilisation solution) was added to each sample, incubated at 95 °C for 2 min, cooled on ice for 2 min then centrifuged at 3000 xg for 2 min. 4.5 μl library preparation enzyme (diluted 3x with H_2_O) was added to each tube, then samples were incubated at 16 °C for 20 min, then 24 °C for 20 min, 37 °C for 20 min and 75 °C for 5 min. Finally, 85 μl amplification master mix (11 μl amplification buffer, 66.5 μl H_2_O, 7.5 μl WGA polymerase) was added to each tube and samples were amplified by PCR (initial denaturation 95 °C for 3 min, then 24 cycles of denaturation 95 °C for 30 s and annealing/extension 65 °C for 5 min).

Amplified DNA was purified using Ampure XP beads. 61.5 μl beads were mixed with 77 μl amplified sample in a fresh 96-well plate and incubated at room temperature for 5 min. The plate was placed on a magnetic stand for 5 min, then the supernatant was discarded and the beads were washed twice with 200 μl freshly prepared 80% ethanol. After the second ethanol wash was discarded, the beads were air dried for 5 min and then resuspended in 45 μl H_2_O and incubated at room temperature for 5 min. The plate was then placed on a magnetic stand and the supernatant transferred to a fresh 96-well plate ready for NGS library prep.

Libraries were prepared using the Illumina Nextera library preparation kit following the manufacturer’s instructions. DNA concentration of the final libraries was determined using a Picogreen fluorescence assay (ThermoFisher) and libraries were pooled at equimolar concentration ready for sequencing on an Illumina NextSeq machine.

### Data processing

Multiplex-GAM sequencing reads were aligned to the mouse mm9 genome assembly using bowtie2, then positive windows were called by GAMtools v0.0.1 using a fixed read threshold of 4. Normalized linkage disequilibrium (D’) matrices at 40 kb genomic resolution were generated by GAMtools (Beagrie and Schueler, 2017).

### SLICE analysis

To convert pair or triplet co-segregation frequencies to interaction probabilities (Pi), we computed the segregation probabilities *v_i_* for a single locus under assumption of spherical shape, with average nuclear radius R (estimated by use of cryosection images as equal to 4.5μm; Beagrie et al., 2017). The co-segregation probabilities *u_i_* for pairs of loci in a not-interacting state have been estimated from GAM segregation data; for interacting loci, instead, we estimated co-segregation *t_i_* probabilities by assuming their physical distance less than slice thickness (*h*≃220 nm). From linear combinations of these probabilities, by use of “mean-field” approximation, we computed the probability of locus segregation in a nuclear profile for pairs (*N_i,j_*) and triplets (*N_i,j,k_*; Supplementary Note).

The expected number of nuclear profiles *M_i_* with 0, 1 or 2 loci, are therefore computed from *N_i,j_* probabilities. From these, in turn, it is also possible estimate the co-segregation ratio *M_1_/(M_1_ +M_2_)*, i.e., the fraction of tubes with two loci on all non-empty tubes. Since the equations describing the tubes content depend on the interaction probability *Pi*, the latter can be estimated by fitting the experimental value of co-segregation ratio (Supplementary Note). The same procedure has been used to estimate the probability of triplet interaction. Significant SLICE contacts are those with a co-segregation ratio greater than the 95^th^ percentile of the expected distribution of co-segregation ratios for two non-interacting loci at the given genomic distance.

To apply SLICE to merged multiplex-GAM dataset, we used a mean field approximation. It consists of introducing a non-integer number of nuclear profiles per tube, obtained as the average of different number of nuclear profiles NPs in the different datasets, weighted with corresponding number of tubes (Supplementary Note).

### Creation of *in silico* merged multiplex-GAM data

Segregation tables (in which each row corresponds to a genomic window, each column to a GAM library and the entries indicate the presence or absence of each window in each NP) were generated from 1NP GAM libraries. A new segregation table was then generated by randomly selecting two, three or four columns from the original table (i.e. 2/3/4x 1NP libraries), combining them into a single column such that the new column is positive if any of the original columns were positive, and removing the columns from the original table. This procedure was performed iteratively until all columns from the original table had been combined. The new, *in silico* combined table was then used for the calculation of normalized linkage disequilibrium matrices.

### SLICE enrichment tests

Enrichment of Active/Enhancer/Inactive/Intergenic windows in pairwise SLICE interactions and analysis of triplet SLICE interactions was carried out as previously described (Beagrie et al., 2017).

### TAD calling

For calling TADs, we aimed to exclude potential effects of using different TAD callers for GAM and Hi-C. Thus, we applied the insulation square method (Crane et al., 2015) to GAM matrices of normalized linkage disequilibrium scores and to Hi-C matrices of normalized ligation frequencies (GSE35156; Dixon et al., 2012), both at 40 kb resolution. We adjusted the insulation square method to also consider negative values from GAM normalized linkage disequilibrium and applied it to contact matrices of each chromosome (im mean, ids 50000, nt 0.1, yb 1.5, bmoe 3). While TAD sizes were nearly insensitive to the size of the insulation square for Hi-C data (reaching plateau around 500 kb square size; Supplementary Figure 5a), increased sizes of the insulation square produced larger TADs for GAM data. Here, we selected a window size of 400 kb for GAM and Hi-C data which maximizes the agreement between both TAD sets as well as to the HMM TAD boundaries published for the Hi-C dataset (Dixon et al., 2012). To consider differences in the number of detected TAD boundaries, we assessed the agreement between two TAD sets by calculating the product of the distances to the respective closest boundary between both sets with either reference.

We used bedtools v2.27.1 (Quinlan AR and Hall IM, 2010) to check if obtained TAD boundaries were touching or overlapping and merged border ranges keeping their maximum boundary score. In comparisons between TAD boundaries from different datasets we considered boundaries to be matching when the distance reported by bedtools closest was at maximum 1 bin.

We checked for abundance of features at the TAD boundaries centred at their boundary midpoints. For a given genomic mark, we analyzed the mean signal within 500 kb around the identified boundary midpoint in windows of 10 kb resolution using bedtools. We estimated the background by randomizing the boundary positions using chromosome-wise circular shifts.

### Generating peak and feature data

We mapped genomic and epigenomic read data to the NCBI Build 37/mm9 reference genome using Bowtie2 v2.1.0 (Langmead et al., 2009). We excluded replicated reads (i.e., identical reads, mapped to the same genomic location) found more often than the 95th percentile of the frequency distribution of each dataset. We obtained peaks using BCP v1.1 (Chen et al, 2012) in transcription factor mode or histone modification mode with default settings. A full list of all features analyzed in this study is given in Supplementary Table 2. We computed mean counts of features for all genomic 40 kb windows by the bedtools window and intersect functions.

### PCA compartments

We computed compartments on GAM and Hi-C data as described (Beagrie et al., 2017; Lieberman-Aiden et al., 2009) or used published compartments definitions (Dixon et al., 2012).

### Identification of differential contacts

GAM and Hi-C implement two different approaches to assess chromatin structure and measure underlying contact frequencies. Comparing the value distributions from GAM D’ and log-scaled Hi-C ligation frequencies highlights differences in the distributions that need to be considered. To directly compare contact intensities between both methods genome-wide in an unbiased fashion, we devised a strategy where we first normalize contact frequency matrices, subtract normalized contact intensities, and, ultimately, define strong contacts seen in both or at significantly different levels by either of the two methods.

In order to avoid amplification of spurious contacts from noise and zero inflation, we limited our analysis to 4Mb genomic distance. From GAM contact data at 40 kb resolution, we removed all contacts with negative D’ values. We also excluded all contacts established between potentially oversampled or undersampled genomic windows. Here, we used the relative number of slices with a positive window (window detection frequency, *wdf*) as a proxy for detectability and removed windows with a *wdf* of less than 5% or above 20%. From the Hi-C data, we excluded all contacts for which zero ligations events were detected. All contacts excluded from any of both datasets were not considered in the definition of differential contacts.

For each chromosome, we applied the z-score transformation to GAM and Hi-C contacts of given distance d. We found this transformation to yield a stable transformation for value intensities from GAM and Hi-C contact matrices (Supplementary Fig. 6c), each resulting in a distribution with a very good fit to a Normal distribution that accounts for the observed decay of mean contact frequency over distance.

To identify the differential contacts between GAM and Hi-C, we computed a delta matrix for each chromosome by subtracting the normalized Hi-C matrix from the normalized GAM matrix. Next, we selected the most differential contacts based on the expected value intensities by fitting a Normal distribution to the distribution of delta values of each chromosome (fitdistrplus R package). We defined Hi-C-specific and GAM-specific contacts to be located within the expected 5% and 95% tails of the fitted distributions, which resulted in sets of 231,165 and 265,157 contacts, respectively. Similarly, we defined a set of strong-and-common contacts which have little difference in the normalized value intensities from GAM and Hi-C. We ranked all contacts with a delta z-score of less than 1.0 according to the lower z-score value from GAM and Hi-C, and extracted the strongest 10% of contacts for each chromosome (total 506,621 contacts).

### Feature enrichments within differential contacts

We queried whether contacts identified to be specific for GAM and Hi-C are associated with specific biological features. First, we checked for the occurrence of homotypic and heterotypic feature combinations in the subsets of Hi-C-specific, GAM-specific, strong-and -common contacts. We annotated 50.29%, 69.58%, and 60.28% of contacts with feature pairs, respectively (Supplementary Fig. 7a).

To assess the amplification of genomic feature pairs in GAM-specific, Hi-C-specific, and strong-and-common contacts, we produced three permutations of random background sets for each of the foreground sets. Here, we randomly sampled the same number of contacts out of all contacts of the same chromosome with the same genomic distance. To determine whether feature combinations are observed in the foreground sets more frequently than to be expected, we computed the amplification (relative frequency) in contrast to the matched background.

Pairs of different features can often be found co-present at the same anchor points of a contact. We applied the UpSetR package (Conway et al., 2017) to the sets of contacts from GAM-specific, Hi-C-specific, strong-and-common, and the genome-wide background and plotted the abundance of a feature pair according to the percentile of feature occurrence along with the number of observed colocalization events between pairs of features. We established the genomic background set by randomly selecting 5% of all non-zero contacts obserbed by GAM and Hi-C.

To directly compare GAM-specific and Hi-C-specific contacts and to see whether both sets can be distinguished using the associated feature pairs, we trained a Random Forest model (sklearn 0.19.2, 100 trees, no pruning) and extracted the top 12 feature pairs that were most informative for binary classification based on the Gini importance criterion.

### Analysis of GAM-preferred and Hi-C-preferred regions

We assessed for each genomic 40 kb window the preference towards preferentially producing GAM-specific contacts or Hi-C-specific contacts. First, we counted how often a window was an anchor point for contacts of the GAM-specific or Hi-C-specific subsets. Next, we calculated the absolute difference between both counts and estimated a 90% percentile cutoff for each chromosome. We considered genomic windows above this threshold to either hold mostly GAM-specific contacts or mostly Hi-C-specific contacts. In total, this resulted in 6,520 windows to be labelled as GAM-preferred regions by having predominantly GAM-specific contacts, and 5,926 Hi-C-preferred regions with a much higher count of Hi-C-specific contacts over GAM-specific contacts. Similarly, we identified genomic regions that are equally well detected by GAM and Hi-C. Here, we counted for each genomic window the number of anchor points from contacts of the strong-and-common set and selected the top 10% genomic windows with the highest counts from each chromosome.

Next, we assessed gene density and transcriptional activity in groups of genomic regions using published gene annotations and mESC-46C TPM values (Ferrai et al., 2017). We transferred provided mm10 gene positions to mm9 using UCSC liftover (Kuhn et al., 2013) and assigned genes to genomic windows of 40 kb using bedtools intersect. We annotated lamina associations within 40 kb genomic regions according to mESC LaminB1 HMM calls (Peric-Hupkes et al., 2010). The genome-wide LAD ratio was computed by the number of positive HMM state calls over the total number of windows. For markers of TFs, histone modifications, and RNA Polymerase II states, we used 40 kb window classification for peak and feature presence (Supplementary Table 2) and counted the number of positive windows in each subset.

### Analysis of interaction complexity

We used SLICE to compute the three-way probability of interaction (*Pi_ABC_*) for all potential 1Mb intrachromosomal triplets from the GAM-1250 dataset (Supplementary Note). We define complexity as the mean *Pi_ABC_* – either the mean over all combinations of “B” and “C” windows for a given “A” window (where complexity is being calculated for a genomic region, Fig. 7c,d) or the mean over all “C” windows for a given “A” and “B” window (where complexity is calculated for a pairwise contact, Fig. 7e).

For each genomic window labelled as Hi-C preferred regions, common regions, or GAM-preferred regions, we checked for the compartment assignment and correlated the outcome with the complexity at the 1Mb genomic window. Similarly, we estimated the complexity of genomic and epigenetic features by subsetting 40 kb genomic windows according to the presence of TFs, histone modifications, and RNA polymerase II states and presenting the complexity of the respective 1Mb genomic window.

## Acknowledgments

The authors thank all lab members and the 4D nucleome consortium for helpful discussions. We thank Warren Winick-Ng for providing illustrations in Fig. 1c,d. We also deeply appreciate the feedback from Teresa Szczepińska on the processing of differential contacts.

AP acknowledges support from the Helmholtz Association (Germany). AP and MN acknowledge support from the National Institutes of Health Common Fund 4D Nucleome Program grant U54DK107977, and the Berlin Institute of Health (BIH). AP acknowledges support from the Deutsche Forschungsgemeinschaft (DFG; German Research Foundation) under Germany’s Excellence Strategy - EXC-2049 - 390688087. MN thanks for support from CINECA ISCRA Grant HP10CYFPS5 and HP10CRTY8P, by computer resources at INFN and Scope at the University of Naples (MN). LW acknowledges the support of Ohio University’s GERB program. RAB is supported by the Wellcome Trust (209181/Z/17/Z).

## Data availability

The original GAM sequencing data (Beagrie et al., 2017) are available from GEO (GSE64881). GAM sequencing data generated for this study are available as a separate accession (GSE166381).

## Code availability

GAM sequencing samples were processed using GAMtools v1.0.0 which is available at https://github.com/pombo-lab/gamtools/releases/tag/v1.0.0. Custom Python code used for data analysis will be made available at https://github.com/pombo-lab/multiplex-gam-2021.

## Author contributions

AP and MN devised the multiplex-GAM strategy;

RAB, AK and RK produced GAM datasets;

RAB, AK and RK optimised the experimental protocol;

RAB performed GAM data processing, QC and *in-silico* merging experiments;

CA, AS, SB, AMC, and MN developed and implemented the updated SLICE model;

CA and AS applied SLICE to find optimal experimental parameters and performed SLICE analysis of GAM data;

RAB performed enrichment analyses for SLICE contact pairs and triplets;

CJT and MS performed TAD boundary analysis;

CJT and AK processed epigenetic and genomic data;

CJT and MS devised pipeline to extract differential contacts;

CJT generated randomized contact sets;

YZ and YL developed and applied the pipeline for feature pair presence and enrichments over background;

CB performed Random Forest discrimination tests;

CB and CJT performed feature pair co-localization tests;

CJT identified and analysed preferred regions;

RAB and CJT assessed complexity of contacts, preferred regions, and feature-positive genomic windows;

AP supervised GAM experiments and bioinformatics analyses;

LW supervised feature enrichment analyses in specific contacts;

MN supervised SLICE development;

AP, RAB, CJT, AK, CA, AS, YZ, CB, LW, and MN contributed to the interpretation of the results;

RAB wrote the first draft of the manuscript; RAB and CJT designed the figures;

RAB, CJT, and AP wrote the manuscript;

All authors provided critical feedback and helped revise the manuscript.

The authors consider RAB, CJT and CA to have contributed equally to this work.

## Materials & Correspondence

Correspondence and material requests should be addressed to

