## Supplementary material for "Multiplex-GAM: genome-wide identification of chromatin contacts yields insights not captured by Hi-C"

### Supplementary Figures

#### Supplementary Figure 1

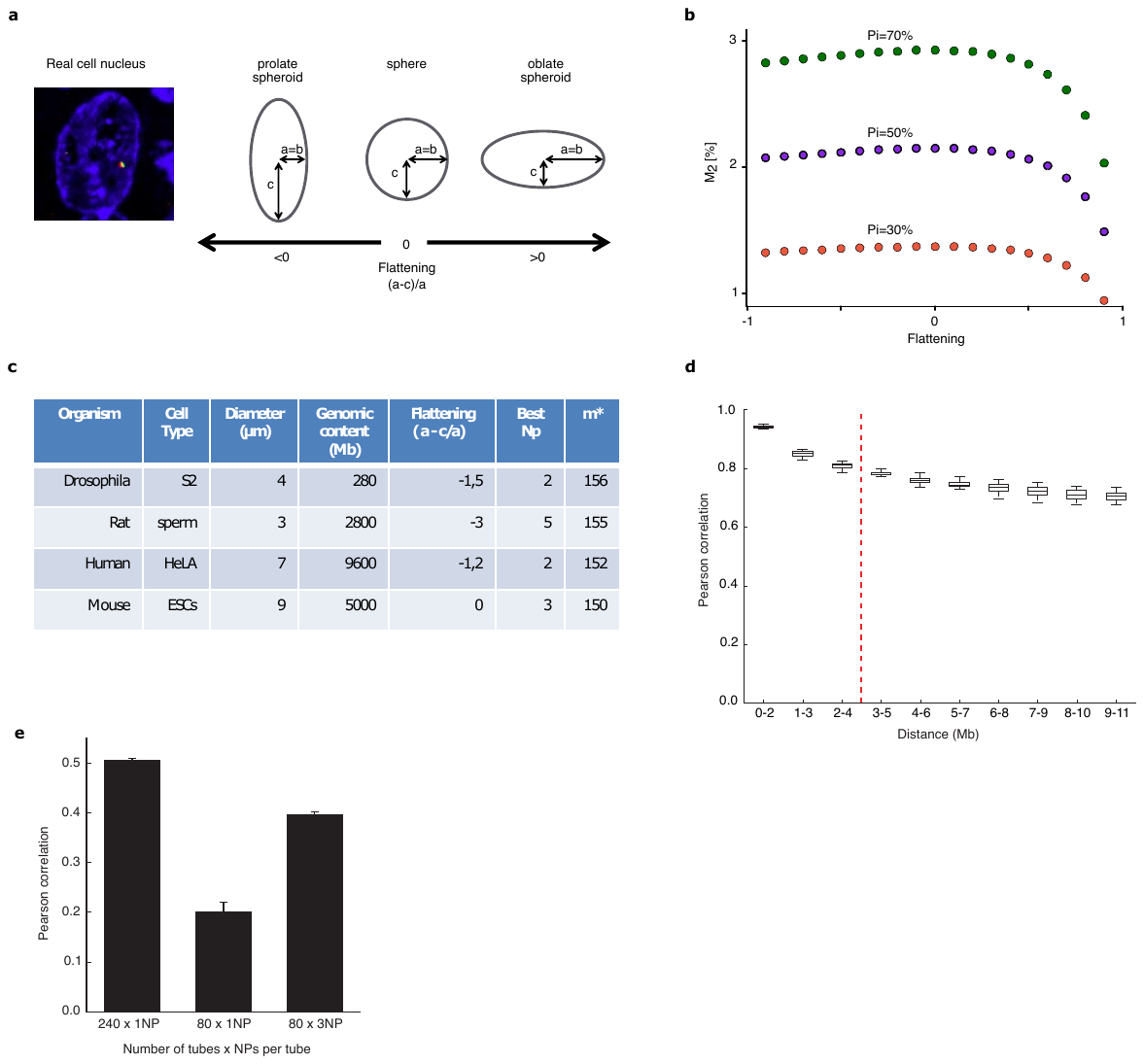

**a**, SLICE can be generalized to account for complex nuclear shapes. As a case study, we considered spheroids (i.e., ellipsoids with two equal axes), which we parametrized by the flattening parameter (the relative difference of the radii). **b**, Co-segregation frequency of pairs of loci (*M2*) derived by SLICE as function of the flattening parameter in spheroidal nuclei of constant volume, for different values of interaction frequency (*Pi*) given slice thickness (*h*) of 220 nm, resolution of 30 kb, detection efficiency of 1 and genomic distances of 50 Mb. **c**, Optimal GAM experimental parameters for different cell types given *h*=220nm, Pi=30% detection efficiency of 1 and genomic distance of 50Mb. The nuclear size and shape of the cell types were obtained from published results (Auger et al.; Brandt et al., 2006; Iborra and Buckle, 2008; Landry et al., 2013). **d**, Correlation of GAM datasets containing single nuclear profiles with GAM datasets where the same NPs are merged *in silico* in sets of three, stratified by distance. **e**, Correlation between two GAM datasets of 240 x 1NP libraries (left) compared to the correlation of two datasets of 80 x 1NP libraries (middle) and two replicates of 80 x 3NP (right, same number of NPs as left bar, same sequencing cost as middle bar). Mean over five randomly sampled subsets ± one standard deviation.

**Supplementary Figure 2**

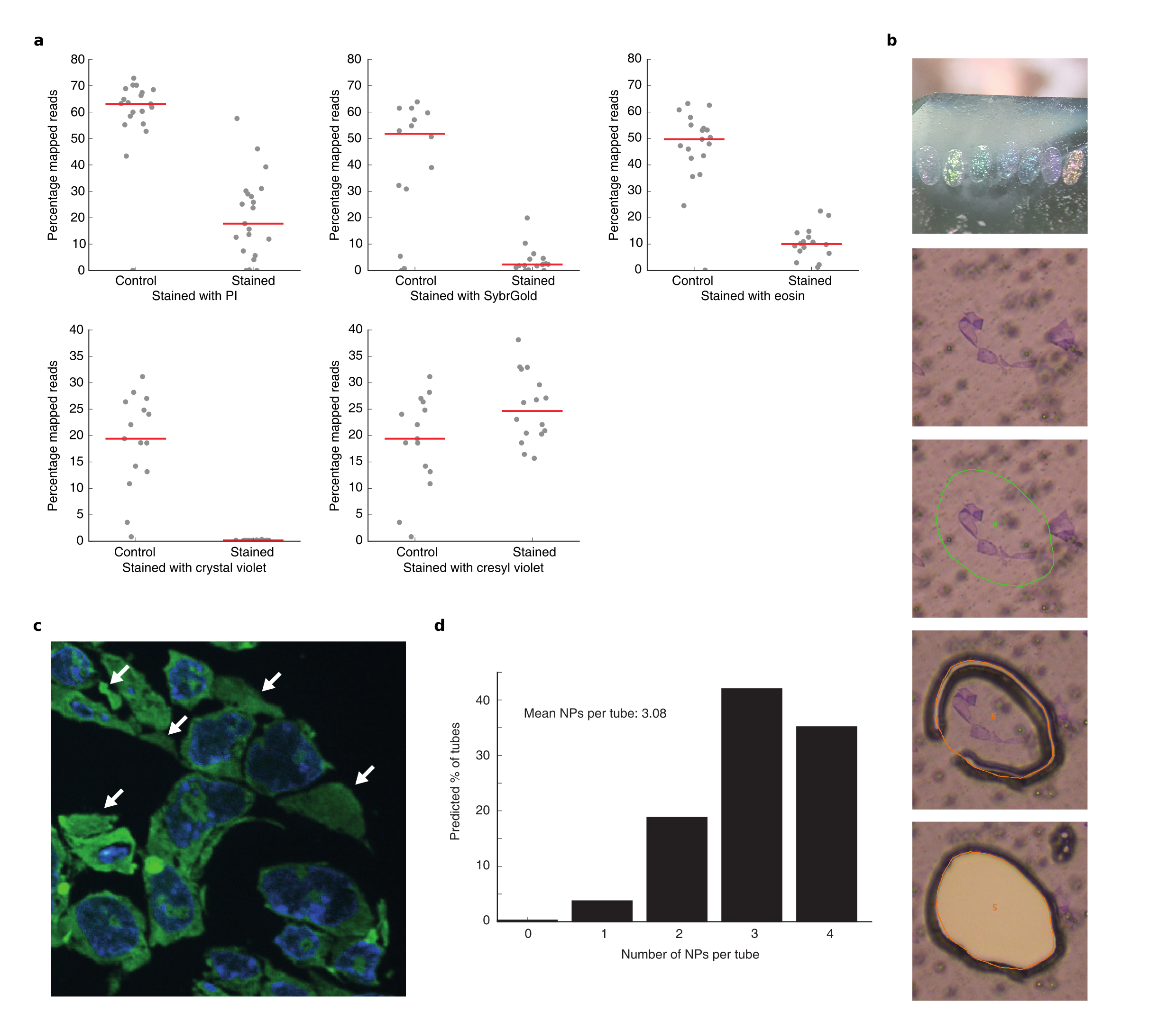

**a**, Effect of various staining protocols on downstream DNA extraction (top row: propidium iodide, SYBR® Gold, Eosin; bottom row: crystal violet, cresyl violet). Red lines indicate the median percentage of mapped reads. **b**, Top panel: cryosection thickness can be identified from the section’s colour. Bottom panels: cresyl violet staining improves the identification of NPs during laser microdissection. **c**, Example mES cell cryosection visualized by confocal fluorescence microscopy, DNA stained with DAPI is shown in blue and RNA stained with SYTO® RNASelectTM dye is shown in green. White arrows indicate cellular profiles that do not intersect nuclei. **d**, Binomial distribution showing the expected number of profiles per sample that intersect nuclei across a collection of samples each with four cellular profiles.

**Supplementary Figure 3**

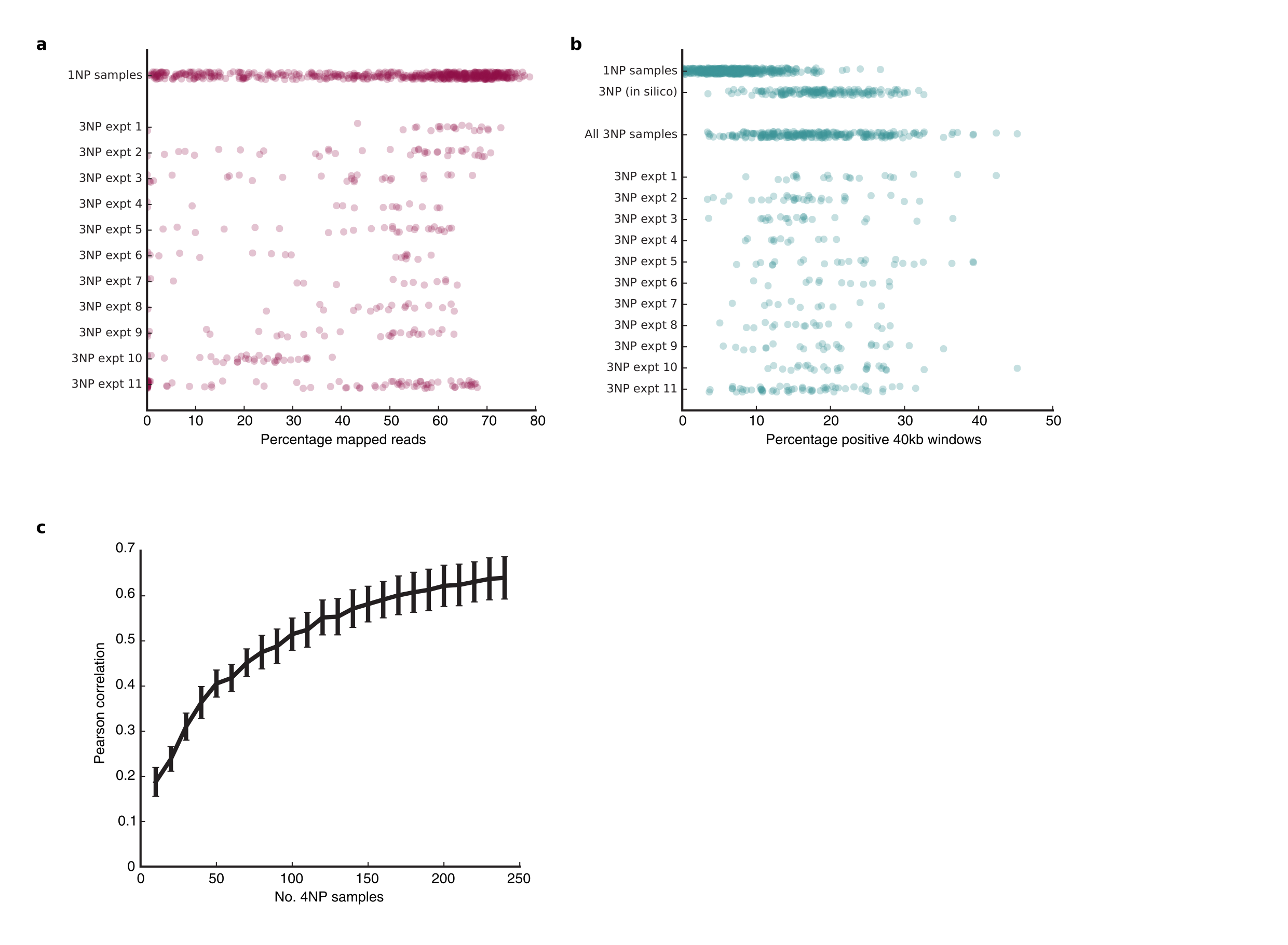

**a**, Percentage of reads mapping to the mouse genome for 1NP libraries and for 3NP libraries separated by experimental batch. **b**, Percentage of positive 40 kb windows or 1NP libraries, for 1NP libraries combined into 3NP libraries *in silico* and for 3NP libraries (i.e. four cellular profiles dissected into each tube with binomial expectation that, on average, three profiles in each tube intersect the nucleus; see Supplementary Fig. 2d,e) separated by experimental batch. **c**, Pearson correlation between normalized linkage matrices calculated from increasing subsets of 3NP libraries and matrix calculated from 481 x 1NP libraries (mean ± standard deviation over mouse autosomes).

**Supplementary Figure 4**

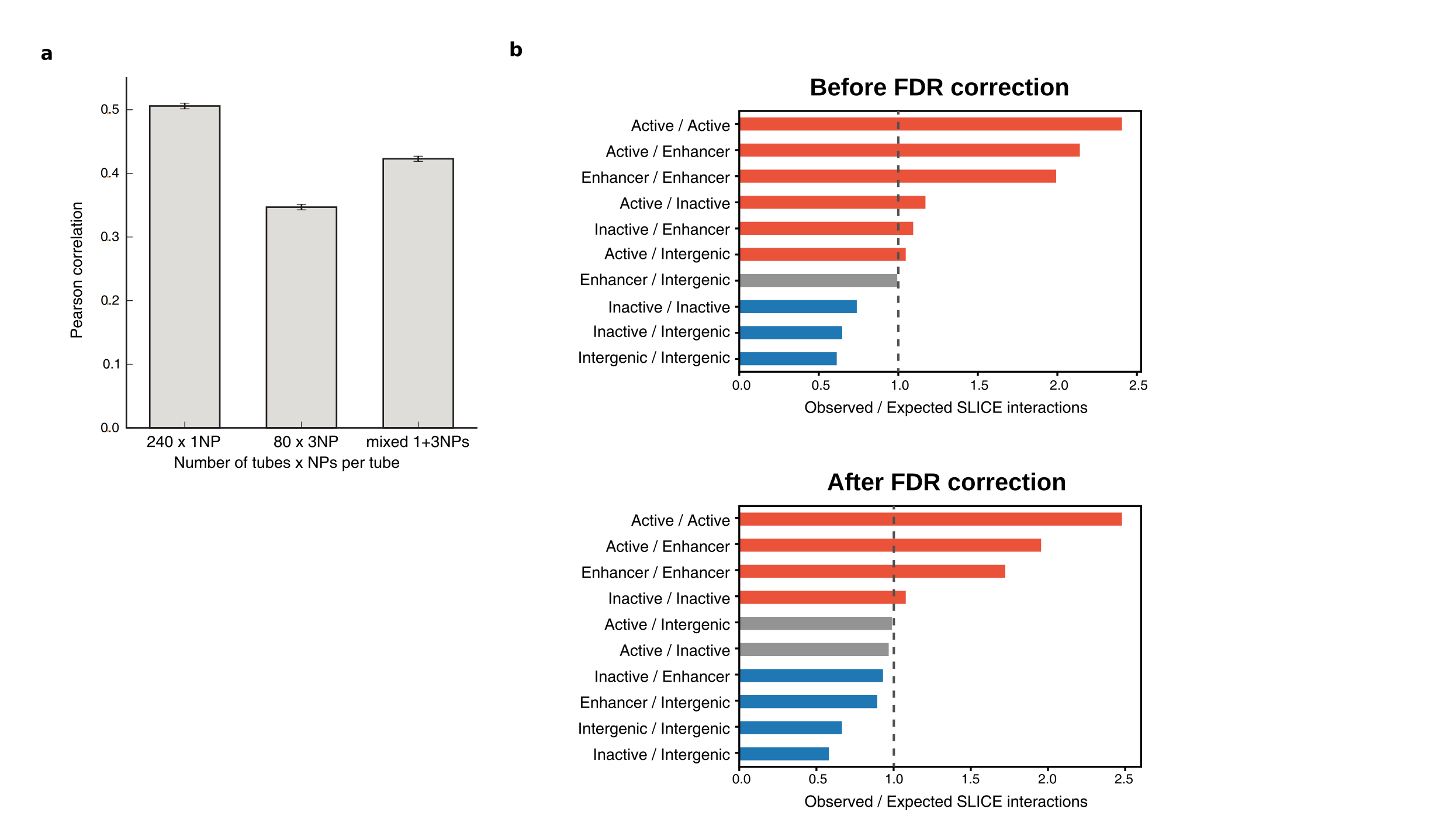

**a**, Correlation between two GAM datasets of 240 x 1NP libraries (left) compared to the correlation of two replicates of 80 x 3NP (middle) and a mixture of 96 x 1NP and 48 x 3NP (right, same number of total NPs mixed in the same ratio as the GAM-1250 dataset). Left and middle bars are the same as in Supplementary Fig. 1e. Mean over five randomly sampled subsets ± one standard deviation. **b**, Enrichment analysis of pairwise interactions identified by SLICE involving active, inactive, intergenic or enhancer regions for the merged GAM-1250 dataset either before (left) or after (right) false discovery rate correction.

**Supplementary Figure 5**

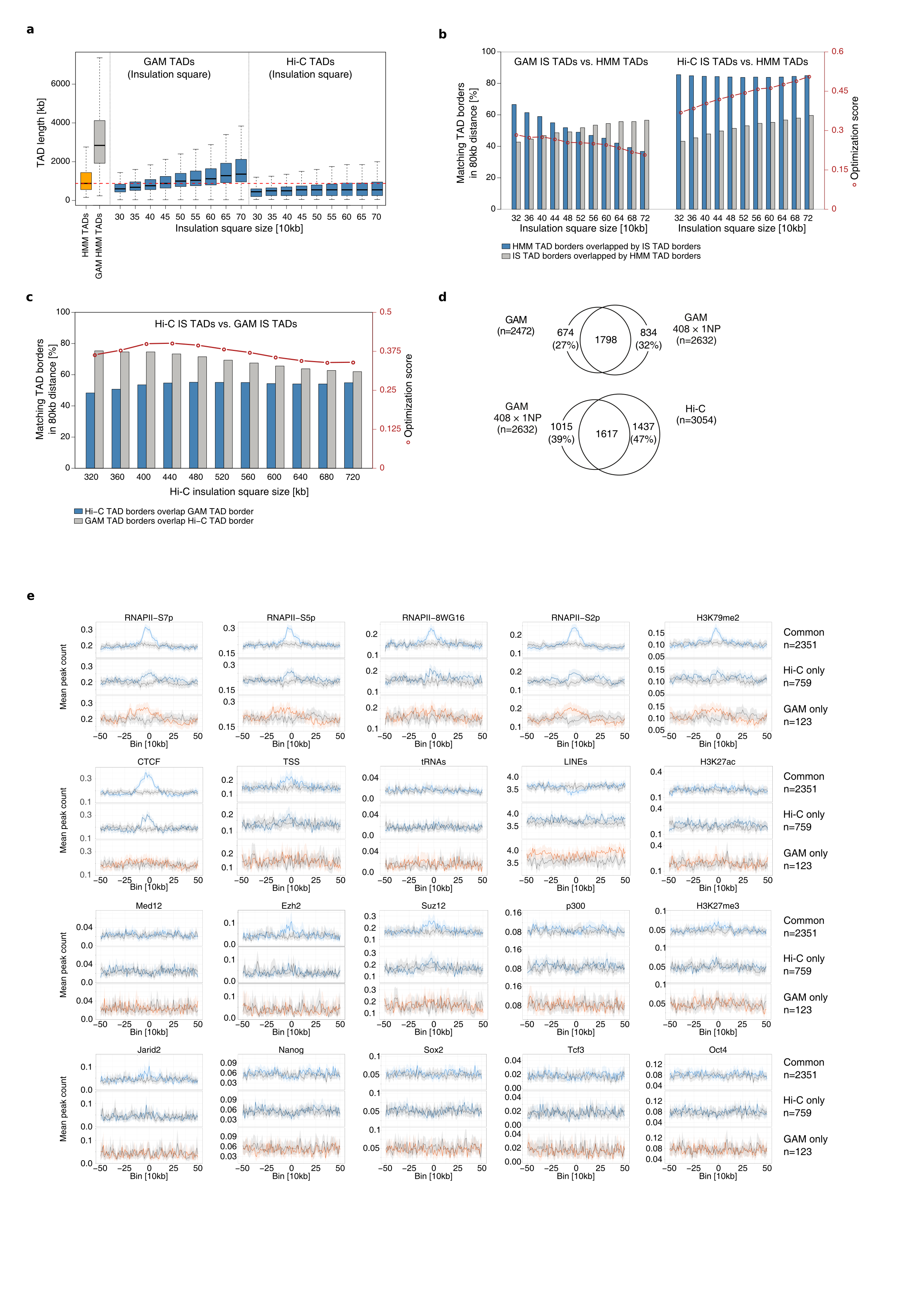

**a**, TAD length for published Hidden Markov Model (HMM) TAD boundaries from mES cells (orange bar), TAD boundaries called from GAM-1250 dataset by HMM (grey bar) or by insulation score (IS) with increasing insulation square size from GAM and Hi-C data at 50 kb resolution. Center lines, median; box limits, upper and lower quartiles; whiskers, 1.5x interquartile range. **b**, Left: Percentage of published Hi-C HMM TAD boundaries overlapping a GAM IS boundary (blue) and percentage of GAM IS boundaries overlapping a Hi-C HMM boundary (grey) for increasing insulation square size. Right: same as left panel but for Hi-C IS boundaries. Optimization score is the product of the two overlap fractions (red line). **c**, Percentage of Hi-C IS TAD boundaries overlapping a GAM IS TAD boundary (blue) or vice versa (grey) with increasing insulation square size. **d**, Top: overlap of IS TAD boundaries calculated from GAM-1250 and the published GAM dataset for mES. Bottom: Overlap of IS TAD boundaries calculated from Hi‑C data or from published GAM dataset. **e,** Aggregated counts for transcription factor binding sites and epigenetic marks centred at TAD boundary sites. Common, Hi-C-specific, GAM-specific boundaries are shown in dark blue, blue, and red, respectively, grey values are based on randomized boundary positions of each category.

**Supplementary Figure 6**

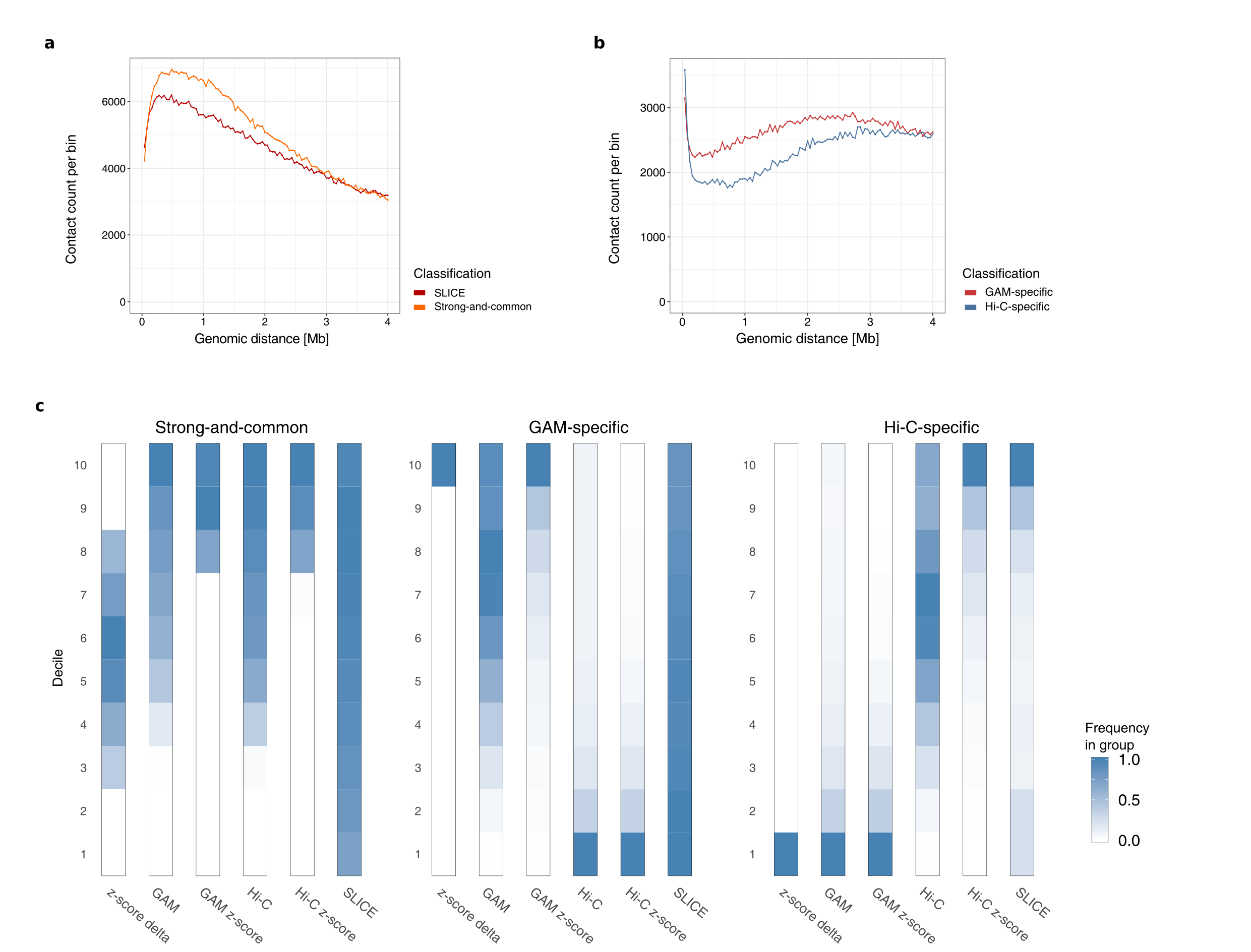

**a**, Distribution by genomic distance of GAM-specific (red) and Hi-C specific (blue) contacts. **b**, distribution by genomic distance of SLICE interactions (red) and strong-and-common contacts (orange). **c**, Decentile distributions of contact intensities before and after z-score normalization for the sets of GAM-specific contacts, Hi-C-specific contacts, and Strong-and-common contacts.

**Supplementary Figure 7**

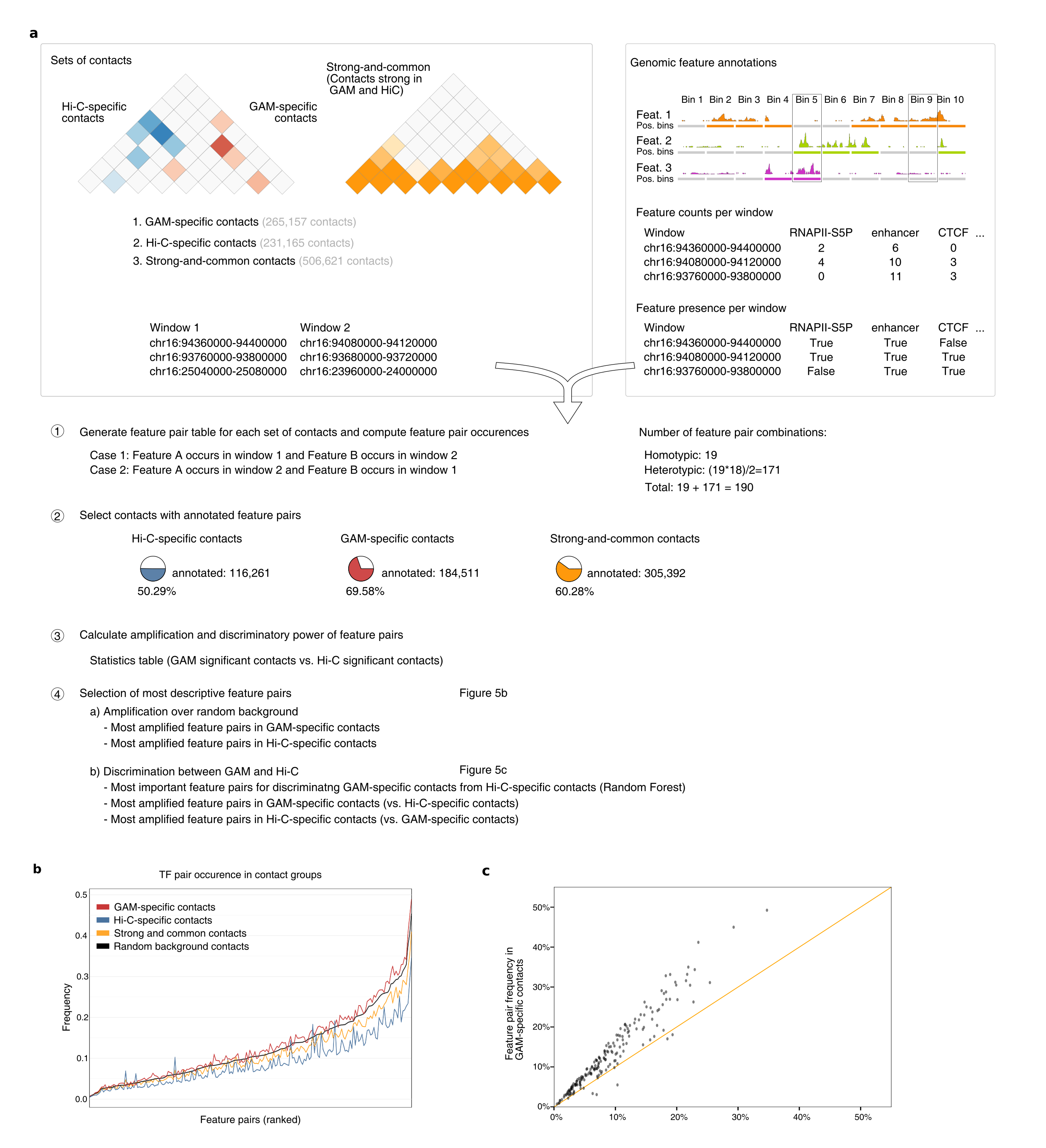

**a**, Strategy for detecting enrichment of feature pairs within sets of specific contacts. **b**, Frequency of feature pair combinations in the sets of GAM-specific contacts, Hi-C-specific contacts, and strong-and-common contacts ordered by the expected presence in the genomic background (5% random sample of contacts). **c**, Correlation of observed frequencies of feature pairs between GAM-specific and Hi-C-specific contacts.

**Supplementary Figure 8**

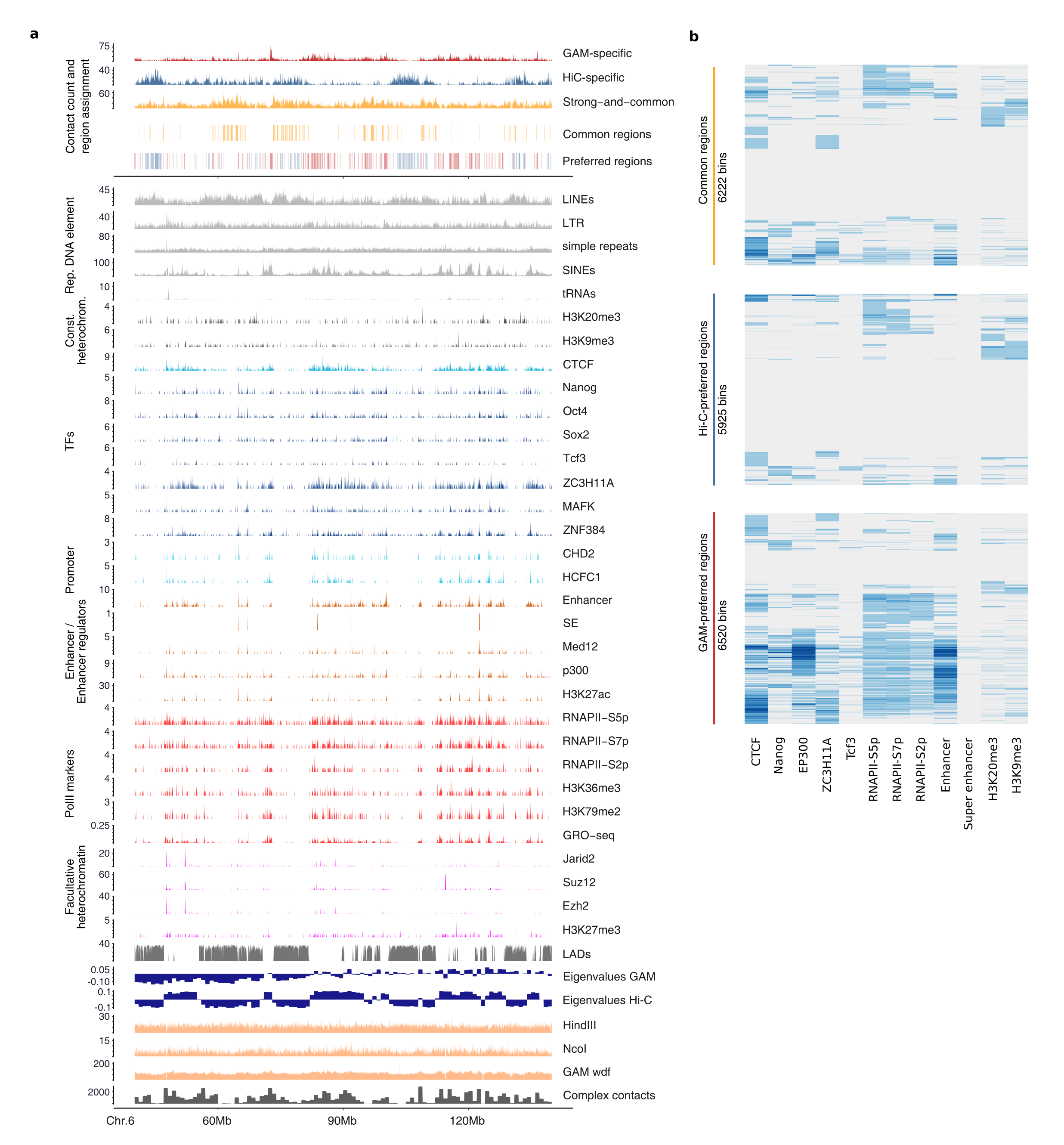

**a**, Feature tracks and assignment of genomic regions having mostly GAM-specific contacts (GAM-preferred regions) or Hi-C-specific contacts (Hi-C-preferred regions). **b**, Heatmap representation for feature presence at GAM-preferred regions, Hi-C-preferred regions) or regions with contacts found at similar intensity by both methods (common regions).

**Supplementary Figure 9**

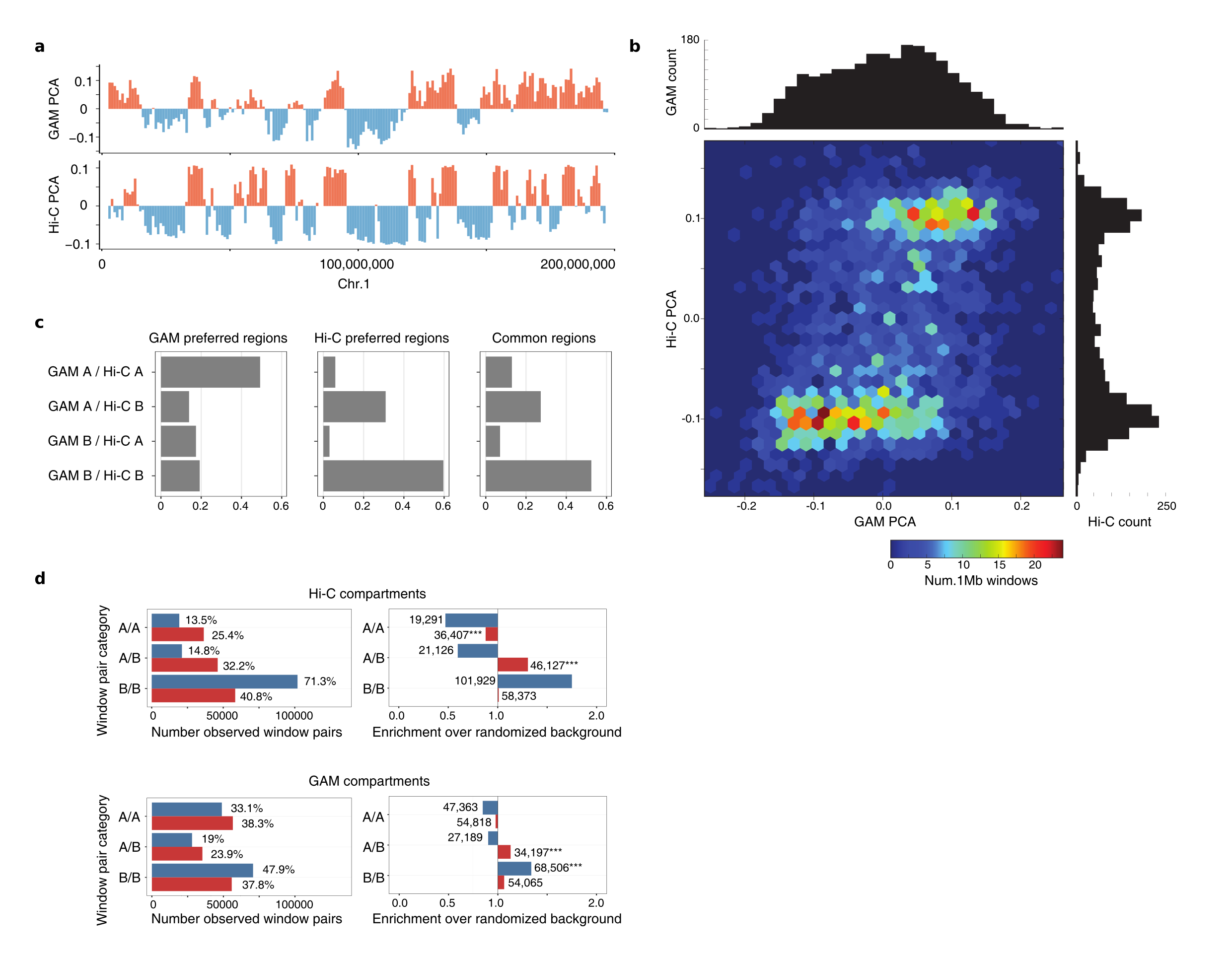

**a**, PCA eigenvalue values from GAM and Hi-C for mouse chromosome 1. **b**, Correlation heatmap for GAM and Hi-C eigenvalues. A small subset of windows assigned to the A compartment by GAM (positive eigenvalues) are assigned to the B compartment by Hi-C (negative eigenvalues). Value histograms indicate that Hi-C eigenvalues are clearly bimodal distributed while GAM finds an increased number of eigenvalues in the transition zone. **c**, Agreement between GAM and Hi-C compartment calls for GAM-preferred (left), Hi-C preferred (middle) and common (right) regions. **d**, Absolute number (left) and enrichment over random background (right) of compartment combinations observed in the sets of Hi-specific contacts (blue) and GAM-specific contacts (red). Compartment definitions of 1Mb resolution were obtained from Hi-C data (top) or GAM data (bottom).

**Supplementary Table 2**

| **Repetitive DNA elements** | |  |  |
| --- | --- | --- | --- |
|  | LINEs, SINEs, LTR, simple repeats, tRNAs | RepeatMasker/UCSC Table Browser | |
| **Constitutive heterochromatin** | | |  |
|  | H3K9me3 | Bilodeau et al., 2009 | GSE18371 |
|  | H3K20me3 | Mikkelsen et al., 2007 | GSE12241 |
| **CTCF** |  |  |  |
|  | CTCF | Shen et al., 2012 | GSE29184 |
| **TFs** |  |  |  |
|  | Nanog | Marson et al., 2008 | GSE11724 |
|  | Oct4 | Marson et al., 2008 | GSE11724 |
|  | Sox2 | Marson et al., 2008 | GSE11724 |
|  | Tcf3 | Marson et al., 2008 | GSE11724 |
|  | ZNF384 | ENCODE |  |
|  | MAFK | ENCODE |  |
|  | ZC3H11A | ENCODE |  |
| **Enhancers** |  |  |  |
|  | Enhancers | Cruz-Molina et al., 2017 | |
|  | Super enhancers | Whyte et al., 2013 | |
| **Enhancer regulators** | |  |  |
|  | Med12 | Kagey et al., 2010 | GSE22557 |
|  | p300 | Shen et al., 2012 | GSE29184 |
|  | H3K27ac | Shen et al., 2012 | GSE29184 |
| **RNA polymerase II activity and transcription-related histone marks** | | | |
|  | GRO-seq | Min et al., 2011 | GSE27037 |
|  | RNAPII-S7p | Ferrai et al., 2017 | GSE94364 |
|  | RNAPII-S5p | Ferrai et al., 2017 | GSE94364 |
|  | H3K4me3 | Shen et al., 2012 | GSE29184 |
|  | RNAPII-S2p | Brookes et al., 2012 | GSE34520 |
|  | H3K36me3 | Marson et al., 2008 | GSE11724 |
|  | H3K79me2 | Marson et al., 2008 | GSE11724 |
| **Facultative heterochromatin** | | |  |
|  | Ezh2 | Peng et al., 2009 | GSE18776 |
|  | H3K27me3 | Ferrai et al., 2017 | GSE94364 |
| **LADs** |  |  |  |
|  | mESC HMM state calls | Peric-Hupkes et al., 2010 | GSE17051 |

**Supplementary Notes**

**for**

**“Multiplex-GAM: genome-wide identification of chromatin**

**contacts yields insights not captured by Hi-C”**

In this Supplementary Note, we will discuss the mathematical implementation of the SLICE (StatisticaL Inference of Co-sEgregation) model and its new features. SLICE allows the quantification of the interaction frequencies between specific DNA loci from GAM (Genome Architecture Mapping) raw data, by correcting for both technical (e.g., detection efficiency) and biological (e.g., genomic distance) factors (Beagrie et al., 2017).

In **Section 1**, we briefly summarize the general mathematical framework of SLICE. **Section 2** is devoted to the extensions of the original SLICE implementation (Beagrie et al., 2017) to model nuclear shape (**Section 2.1**), cellular ploidy (**Section 2.2**) and the multiplex-GAM experimental setup (**Section 2.3**). Finally, we describe how SLICE can be employed to optimize GAM experimental protocols (**Section 3**) and to combine different GAM datasets (**Section 4**).

1. **SLICE model**

A GAM experiment consists of the collection of many nuclear cryosections (nuclear profiles or NPs) in random orientations from a population of nuclei, with one NP being isolated from each nucleus. The DNA content of each NP is then sequenced to provide information about presence/absence of any DNA locus. SLICE is a statistical model that, starting from raw GAM data, estimates the interaction probability between two or more chromatin loci.

In the following, we briefly introduce the SLICE model focusing first on the case of pairwise (**Section 1.1**) and threewise (**Section 1.2**) interactions. For additional details, we refer the reader to Beagrie et al., 2017.

**1.1	Pairwise interactions**

In the SLICE model, we assume that a pair of loci, *A* and *B*, can be in two different states: in an interacting state, when the two loci are spatially close even if genomically distant, or in a non-interacting state, when they are separated from each other with a physical nuclear distance that is dependent on their genomic separation. These states occur with probabilities *Pi* and *(1-Pi)* respectively. In addition, we define state-specific probabilities for the loci *A* and *B* to be segregated in a randomly selected slice (**Suppl. Notes Figure SN1**) as follows:

- v_1_ and v_0_ are the probabilities to find/not find a single locus in the slice;
- u_2_, u_1_, u_0_ are the probabilities to find 2, 1 or 0 loci in a pair of non-interacting loci;
- t_2_, t_1_, t_0_ are the probabilities to find 2, 1 or 0 loci in a pair of interacting loci.

The following normalization relationships are satisfied:

- u_2_ + 2u_1_ + u_0_ = 1
- t_2_ + 2t_1_ + t_0_ = 1
- v_1_ + v_0_ = 1

These probabilities depend on the geometric properties of the system. For instance, in the case of spherical nuclei, the probabilities v_0_ and v_1_ can be analytically calculated and depend on the ratio between the slice thickness and the nuclear radius (Beagrie et al., 2017). Analogously, the probabilities for pairs of loci (i.e. t_0_, t_1_, t_2_ and u_0_, u_1_, u_2_) depend also on the average physical distance between the loci in the interacting and non-interacting states, and can be estimated numerically (e.g., via Monte Carlo simulations) or from GAM data (Beagrie et al., 2017). However, for loci at large genomic distances ($\gtrsim$50Mb) and small physical distances *d_I_* when interacting (*d_I_* $\lesssim$ *h*), the following approximations can be used (see Beagrie et al., 2017):

$$u_{0}\sim v_{0}^{2}, u_{1}\sim v_{1}v_{0}, u_{2}\sim v_{1}^{2}$$

$$t_{0}\sim v_{0}, t_{1}\sim0, t_{2}\sim v_{1}$$

From these probabilities, SLICE can predict the outcome of a GAM experiment as a function of the interaction probability *Pi* of a pair of loci. For example, if only one nuclear profile is added per tube, the fraction of tubes that include neither *A* nor *B* loci in a diploid nucleus is:

*M_0_* =*N_0,0_*= [*Pi* t_0_ + (1-*Pi*)u_0_]^2^

where *N_0,0_* is the fraction of slices with neither *A* nor *B* loci.

Similarly, we can calculate the fraction of tubes including two (*M_2_*) or one (*M_1_*) loci (Beagrie et al., 2017). Conversely, *Pi* can then be estimated by comparing the predictions of *M_2_*, *M_1_*, *M_0_* with the experimental values measured by GAM (Beagrie et al., 2017).

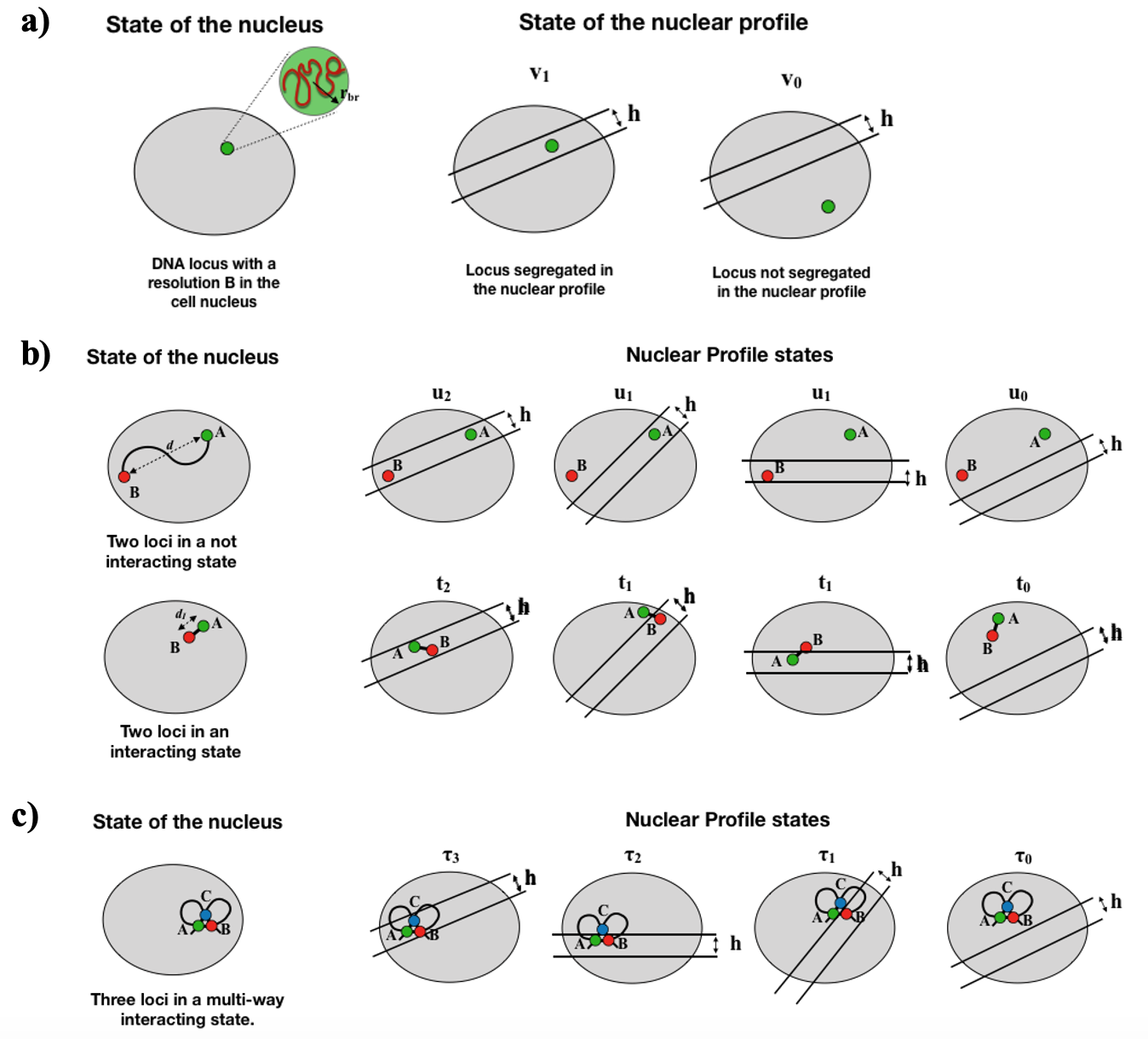

**Figure SN1**: SLICE model parameters.

a) Schematic picture of a DNA locus in the cell nucleus with a given resolution *b_r_*. Each locus is represented by a sphere with a radius $r_{b_{r}}$*,* that depends on the considered resolution *b_r_.* The probabilities to find or not a locus in a population nuclear profiles, with slice thickness *h*, are v_1_ and v_0_ respectively.
b) The probability to find a pair of loci in a population of nuclear profiles depends on whether they are in a not-interacting (u_i_, top) or interacting (t_i_, bottom) state.
c) Probability τ_i_ of finding 3, 2, 1 or 0 loci of a triplet in a interacting state.

**1.2	Threewise (triplet) interactions**

SLICE can exploit the co-segregation information from GAM to estimate the frequency of multivalent interactions between DNA loci. For example, the interaction probability of triplets of loci, *Pi_3_*, can be estimated following an approach similar to the pair-wise calculation. Briefly, SLICE can estimate how often any combination of the loci in the triplet are co-segregated in the same nuclear slice in the case when the three loci are interacting or not interacting.

As an example, we consider the case where all the three loci *A, B* and *C* are interacting and define τ_i_ the probability that i (i = 0, 1, 2, 3) loci of the triplet are co-segregated in the same nuclear profile. Again, these probabilities can be numerically estimated. However, if the physical distances of the loci interacting in the triplet is $\lesssim$*h*, *h* being the slice thickness, the probabilities τ_i_ are well approximated by:

τ_0_ ~ v_0_,	τ_1_ ~ τ_2_ ~ 0, τ_3_ ~ v_1_

where v_1_ and v_0_ have been previously defined.

Analogously, SLICE can calculate the probabilities corresponding to the cases where only two loci of the triplet are interacting or the three loci are not interacting (Beagrie et al, 2017). These quantities can be used to derive the equations for the fraction of nuclear profiles *N_i,j,k_* (with i, j, k = 0, 1) containing any combination of the loci. Hence, the fraction of tubes containing any number of loci, which are the quantities measured by GAM and can be fitted to determine the triplet interaction probability (Beagrie et al., 2017).

In those calculations, additional complexities can be taken into account, such as the detection efficiency of loci in a tube or the effects of genomic resolution (as discussed in Beagrie et al, 2017).

2. **SLICE model extensions**

In its previous implementation, SLICE considered diploid cells with nuclei approximated as spheres (Beagrie et al., 2017). Here, we show how the model can be generalized to include different nuclear shapes (**Section 2.1**) and ploidies (**Section 2.2**). Finally, in **Section 2.3** we extend SLICE to model the new multiplex-GAM protocol, which reduces hands-on time and costs.

2.1 **Modelling the nuclear shape**

SLICE can model an arbitrary nuclear shape by extending the computation of the probabilities v_i_, u_i_ and t_i_ defined above. In general, these can be numerically estimated (e.g., via Monte Carlo methods). For the sake of simplicity, we consider ellipsoids, which are good approximations of nuclear shapes for a number of cell types (**Suppl. Figure S1c**) and can be treated analytically.

Below, we calculate the analytical expression of v_1_ in ellipsoidal nuclei, which allows the estimation of the probabilities u_i_ and t_i_ for pairs of loci at large genomic distances ($\gtrsim$50Mb) and small physical distances *d_I_* when interacting (*d_I_* $\lesssim$ *h*; see above); for all other cases, numerical simulations or GAM data can be used, as described in (Beagrie et al, 2017). The results can be easily generalized to take into account different experimental detection efficiencies or resolutions (Beagrie et al., 2017).

2.1.1 Segregation probability v_1_ for ellipsoidal nuclei

We consider a cell nucleus with an ellipsoidal shape and semi-axes *a, b, c*. Without loss of generality, any ellipsoid can be described by the canonical implicit equation:

$\frac{{\text{x}_{\text{1}}}^{\text{2}}}{\text{a}^{\text{2}}}\text{ + }\frac{{\text{x}_{\text{2}}}^{\text{2}}}{\text{b}^{\text{2}}}\text{ + }\frac{{\text{x}_{\text{3}}}^{\text{2}}}{\text{c}^{\text{2}}}\text{ = 1}$	(Eq. 1)

where x_1_, x_2_ and x_3_ are the 3D coordinates.

In a coordinate system defined by the following anisotropic scaling transformation:

$\left( \text{x}_{\text{1}}\text{,}\text{ }\text{x}_{\text{2}}\text{,}\text{ }\text{x}_{\text{3}} \right)\text{ → }\left( \text{ x'}_{\text{1}}\text{=}\frac{\text{x}_{\text{1}}}{\text{a}}\text{,}\text{ }\text{x'}_{\text{2}}\text{=}\frac{\text{x}_{\text{2}}}{\text{b}}\text{,}\text{ }\text{x'}_{\text{3}}\text{=}\frac{\text{x}_{\text{3}}}{\text{c}} \right)$	(Eq. 2)

the ellipsoid is transformed into a sphere with unit radius. Hence, in this new coordinate system (indicated by apex symbols) we can use the equations for v_1_ and v_0_ written in the spherical case (see Beagrie et al., 2017), with a slice thickness h’ defined by:

$\text{h' = }\frac{\text{h}}{\sqrt{{\text{(a}\sin\text{ϕ}\cos\text{θ}\text{)}}^{\text{2}}\text{ + }{\text{(b}\sin\text{ϕ}\sin\text{θ}\text{)}}^{\text{2}}\text{ + }{\text{(c}\cos\text{ϕ}\text{)}}^{\text{2}}}}$	(Eq. 3)

where *θ* and ϕ are respectively the azimuthal and polar angle identified by the direction $\hat{n}$ of the slice plane (**Suppl. Notes Figure SN2a**). The average segregation probability for a single locus, v_1_’, can be obtained by integrating over all possible slice orientations and positions, as follows:

$\text{v}_{\text{1}}\text{' = }\frac{\int_{\text{S}}^{\text{ }} \text{d}\text{Ω}\int_{\text{-h'(}\text{θ}\text{, }\text{ϕ}\text{)/2}}^{\text{1}} \text{ }\text{dz}\text{' }\text{v}_{\text{1}}\text{'(z',}\hat{n}\text{)}}{\int_{\text{S}}^{\text{ }} \text{d}\text{Ω}\int_{\text{-h'(}\text{θ}\text{, }\text{ϕ}\text{)/2}}^{\text{1}} \text{dz}\text{'}}$	(Eq. 4)

where $\int_{\text{S}}^{\text{ }} \text{d}\text{Ω}\text{ = }\int_{\text{0}}^{\text{2}\text{π}} \text{dθ}\int_{\text{0}}^{\text{π}} \text{d}\text{ϕ}\text{ }\sin\text{ϕ}\text{ }$ is the integral over the solid angle, $\text{v}_{\text{1}}\text{'(z,}\hat{\text{n}}\text{)}$ is the segregation probability of a single locus in a slice of a sphere with orientation $\hat{\text{n}}$ and distance from the equatorial plane equal to $z$. Since the transformation (**Eq. 2**) is linear, the associated Jacobian determinant is constant. Therefore the volumes ratios are invariant, and then v_1_’ is equal to v_1_.

As an example, in **Suppl. Notes Figure SN2b** we show the value of v_1_ computed by numerical integration of equation (**Eq. 4**) for different values of semi-axis lengths (a, b, c), as a function of the slice thickness *h*. The values computed from **Eq. 4** match those obtained by Monte Carlo simulations (marked by circles).

Given the above equation for v_1_, we can estimate the probabilities u_i_ and t_i_ and the fraction of tubes with 2, 1 or 0 loci (*M_2_*, *M_1_*, *M_0_*) in the case of an ellipsoidal nucleus, with the calculations mentioned in **Section 1** and explained in detail in Beagrie et al., 2017.

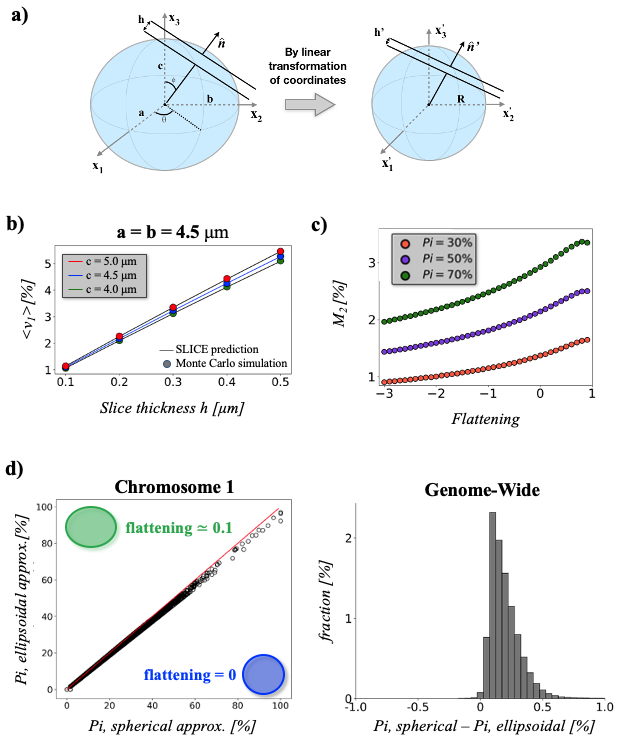

**Figure SN2**: SLICE generalization to ellipsoidal nuclei.

a) Schematic picture showing the linear transformation of coordinates, from (x_1_, x_2_, x_3_) to (x_1_’, x_2_’, x_3_’), for mapping a generic ellipsoid with semi-axis length (a, b, c) to spherical case, with unitary radius (*R*=1).
b) Average probability v_1_ that a locus is found in a nuclear profile from a population of ellipsoidal nuclei, as a function of the slice thickness *h* and for different lengths of semi-axes. Here, we show the spherical case (a = b = c = 4.5μm) and the cases where we increase/decrease the semi-axis length by about 10%. As expected, we find that v_1_ decreases (increases) if we consider a smaller (bigger) nuclear volume. The SLICE prediction (solid lines) matches with independent Monte Carlo simulations (circles).
c) Co-segregation frequency *M_2_* as a function of the nucleus *flattening* parameter*,* for different values of interaction probability *Pi*. Unlike **Suppl.** **Figure S1b**, here we considered variations in semi-axis lengths leading to changes of the ellipsoid volume *V*. We fixed slice thickness h=*220 nm*, *b_r_=30 kb* and detection efficiency *ε=1* and we considered two loci on different chromosomes, and therefore randomly distributed within the nucleus.
d) Comparison of SLICE results on multiplex-GAM data at 40 *kb* resolution, in spherical (*R=*4.5 μm) and ellipsoidal approximation, where we increased the length of one semi-axis of about 10% (a = b = 4.5, c = 5.0 μm; *flattening*≃0.1). (Left) Scatterplot of interaction frequencies *Pi* for chromosome 1 in spherical and ellipsoidal approximation. (Right) Distribution of differences in *Pi* values genome-wide between spherical and elliptical case. For sake of simplicity we only show differences between non-zero *Pi* values in both cases.
2.1.2 Finite genomic resolution of loci

In **Eq.4,** we assumed that the genomic region we want to detect is a dimensionless point. However, to take into account for the finite size of a locus at a given genomic resolution, we can represent a DNA locus including $b_{r}$bases as a sphere with a finite radius $r_{b_{r}}$ (**Suppl. Notes Figure SN1a**). It can be shown that, if $r_{b_{r}}$ << a, b, c (i.e., the semi-axes of the ellipsoidal nucleus), the previous equations still hold if *h* is replaced by an effective slice thickness *h_eff_* = *h* + 2$r_{b_{r}}$ (Beagrie et al., 2017). An estimation of $r_{b_{r}}$ can be obtained under the assumption that the genetic material is uniformly distributed within the cell nucleus, which leads to:

$\text{ }r_{b_{r}} \text{=}{\text{ }\left( \frac{b_{r}}{L}\text{ abc} \right)}^{\text{1/3}}$	(Eq. 5)

where *L* is the total length of the genome (Beagrie et al., 2017).

2.1.3 Effects of nuclear shapes

In this section, we use the equations for ellipsoidal nuclei to investigate how the nuclear shape can affect the co-segregation frequency *M_2_*, i.e. the expected fraction of tubes containing both loci of a pair (**Section 1**). Moreover, we test how robust are the SLICE calculations obtained in mES cells to changes of the nuclear shapes.

While the above equations hold true for an arbitrary ellipsoid, here, for the sake of simplicity, we focus on spheroids, i.e., ellipsoids with two equal semi-axes (a = b) (**Suppl.** **Figure S1a**). In this way. we can parametrize the nuclear shape by using the *flattening*, defined as 1 - c/a*.* By varying the length of the c-axis, we can consider two possible cases: the a and b semi-axes are kept constant, leading therefore to a variation of the total volume, or, alternatively, the volume is kept constant, and the two semi-axes a and b change accordingly. For both cases. we derived the co-segregation frequency of pairs of loci (*M_2_*) as a function of the *flattening* parameter (shown in **Suppl.** **Figure S1b** and **Suppl. Notes Figure SN2c** respectively). In general, we found that *M_2_* changes non-trivially with *flattening* and exhibits a different behaviour with respect to volume variation, proving that not only the nucleus shape, but also its size affects the co-segregation frequency.

Next, we quantified how robust are our SLICE results obtained under the assumption of perfectly spherical nucleus. To this aim, we compared the results obtained on combined Multiplex-GAM data at 40 kb resolution in spherical approximation against the results obtained by introducing a small flattening (a = b = *R* = 4.5 *μm*, c = 5 *μm*, *flattening* ≈ 0.1). In both cases, SLICE finds very similar results (as shown in **Suppl. Notes Figure SN2d**). In particular, this change in the shape leads to a slight shift of interaction probability *Pi*, as quantified by Pearson correlation coefficients (values above the 99%) and overlaps of contacts (about 98% for all the chromosomes). Additionally, the estimated detection efficiencies ε are very similar (88% ellipsoidal case against 86% in spherical case).

2.2 **Modelling cellular ploidy and detection efficiency**

To extend SLICE to cells with any chromosome copy number (ploidy), we generalized the equations for the *N_i,j_*, i.e., the expected fraction of nuclear profiles including *i* copies of locus *A* and *j* copies of locus *B* (see above) in a population of *N* cells. In the following, for the sake of simplicity, we focus on the cases of haploid, diploid and triploid cells, but SLICE equations can be easily generalized to arbitrary ploidies.

For diploid cells, where *i* and *j* assume values between 0 and 2, *N_i,j_* can be written in mean field approximation as a function of c_0_ = *Pi*⋅t_0_+(1-*Pi*)⋅u_0_ and v_0_ , i.e., the probabilities that a pair of loci or a single locus are not contained in a nuclear profile, respectively (Beagrie et al., 2017). More specifically, the *N_i,j_* are polynomial functions of v_0_ and c_0_ with a degree equal to the ploidy of the cells.

For example, in haploid cells the expressions for *N_i,j_* are linear combinations of v_0_ and c_0_:

*N_0,0_* = c_0_

*N_1,0_* = *N_0,1_ =* (v_0_ *-* c_0_)	(Eq. 6)

*N_1,1_* = (*1 - 2*v_0_ *+* c_0_)

On the other hand, for triploid cells the *N_i,j_* are third degree polynomial functions in v_0_ and c_0_. For instance, *N_1,1_*, i.e., the fraction of nuclear profiles including one locus *A* and one locus *B* in a triploid cell is:

*N_1,1_ = 3 (1 - 2*v_0_ *+* c_0_*)* c_0_*^2^ + 6 (*v_0_*-*c_0_*)^2^*c_0_	(Eq. 7)

SLICE can also model experimental limitations by introducing a detection efficiency *ε* defined as the probability that a given locus present in a slice is detected. The expected fraction of nuclear profiles with an *ε* < 1, *N_i,j_ ^ε^*, is a linear combination of the *N_i,j_* values corresponding to *ε* = 1 (Beagrie et al., 2017). As an example, equation **Eq. 7** including a detection efficiency *ε,* becomes*:*

*N_1,1_^ε^ = ε^2^ [N_1,1_ +(1- ε) 2N_2,1_ +(1- ε)^2^ (2N_3,1_ + N_2,2_) +(1- ε)^3^ 2N_3,2_ +(1- ε)^4^ 2N_3,3_]*

**2.3	SLICE for multiplex-GAM datasets**

In this section, we show how SLICE can be generalized for multiplex-GAM. In this new GAM protocol, multiple nuclear profiles are collected into a single sequencing tube. This leads to a significant reduction in reagent costs and labor time and a greater sensitivity (as we show in **Section 3**).

For any pair of loci, the previous equations for the single slices *N_i,j_* still hold true in multiplex-GAM, while the equations for the expected number of tubes with 0, 1 and 2 loci (*M_0_*, *M_1_* and *M_2_*) need to be modified to account for the number of nuclear profiles *X_NP_* in each tube. By indicating with *n* the ploidy, we have:

*M_0_* = (*N*_0,0_)*^XNP^*

*M_1_* = 2 [($\sum_{i =0}^{n} N_{i,0}$)*^XNP^* - (*N_0,0_*)*^XNP^*]	(Eq. 8)

*M_2_* = 1 – *M_1_* - *M_0_*

Similar equations can be written for an arbitrary number of loci. For instance, in the case of triplets (*A*, *B* and *C* loci), we can first estimate the expected fraction of nuclear profiles, *N_i,j,k_* with a number *i*, *j*, *k* of loci *A*, *B*, C. From these, by analogous calculations, we can derive the expected fraction of tubes with 0,1,2 or 3 loci of the triplet (*M_0_*, *M_1_*, *M_2_* and *M_3_*, respectively), in the general case of a number *X_NP_* of nuclear profiles per each sequencing tube:

*M_0_* = (*N*_0,0,0_)*^XNP^*

*M_1_* = [($\sum_{\text{i}\text{ =0}}^{\text{n}} \text{N}_{\text{i,0}\text{,0}}$)*^XNP^* + ($\sum_{\text{j =0}}^{\text{n}} \text{N}_{\text{0,j}\text{,0}}$)*^XNP^* +($\sum_{\text{k =0}}^{\text{n}} \text{N}_{\text{0,0}\text{,k}}$)*^XNP^* - 3(*N_0,0,0_*)*^XNP^*]

*M_2_* = [($\sum_{\text{i}\text{, j}\text{ =0}}^{\text{n}} \text{N}_{i\text{,j}\text{,0}}$)*^XNP^* +($\sum_{\text{i}\text{, k}\text{ =0}}^{\text{n}} \text{N}_{i\text{,}\text{0}\text{,k}}$)*^XNP^* +($\sum_{\text{j,k}\text{=0}}^{\text{n}} \text{N}_{0\text{,j}\text{,k}}$)*^XNP^* -	(Eq. 9)

- 2(($\sum_{\text{i}\text{ =0}}^{\text{n}} \text{N}_{i\text{,0}\text{,0}}$)*^XNP^*+$\sum_{\text{j}\text{ =0}}^{\text{n}} \text{N}_{0\text{,j}\text{,0}}$)*^XNP^*+$\sum_{\text{k =0}}^{\text{n}} \text{N}_{0\text{,}\text{0}\text{,k}}$)*^XNP^*) + 3*N_0,0,0_^XNP^*]

*M_3_* = 1 – *M_2_* – *M_1_* - *M_0_.*

3. **Using SLICE to optimize GAM experimental design**

In this Section, we show how GAM sensitivity can be estimated by SLICE and how this can be used to optimize the GAM experimental parameters (e.g., the number of tubes to sequence, the slice thickness and the number of nuclear profiles per tube, etc.) for a given organism and cell type.

3.1 **GAM sensitivity**

GAM sensitivity in the analysis of pairwise interactions can be defined as the probability to reject the null hypothesis *H*_0_ that a pair of loci is non-interacting (*Pi*=0), when the alternative hypothesis *H*_1_ is true, i.e. when the pair is interacting with a probability *Pi*>0.

As discussed in Beagrie et al., 2017, the interaction probability *Pi* can be estimated from GAM data by fitting the value of the co-segregation probability, i.e., *M*_2_*/*(*M*_1_ + *M*_2_), which, for loci at a given genomic distance, increases with increasing *Pi*. Beyond the expected value of the co-segregation ratio, its full probability distribution can be also estimated from SLICE (see **Suppl. Notes Figure SN3a**).

By using the co-segregation ratio to estimate *Pi*, the GAM sensitivity *Pr* will depend on how distinct the probability distributions of the co-segregation ratio are for *Pi* = 0 and *Pi* > 0. Specifically, for any given combination of experimental parameters and interaction probability *Pi*, we can define *Pr* as the probability to obtain a co-segregation ratio *M_2_*/(*M_2_*+*M_1_*) greater than the 95^th^ percentile of the distribution of co-segregation ratio at *Pi* = 0 (Beagrie et al., 2017, **Suppl. Notes Figure SN3a**).

Analogously, it is also possible to define GAM sensitivity to detect triplet interactions. Here, the co-segregation ratio is defined as *M_3_*/(*M_1_* + *M_2_* + *M_3_*). Then, the sensitivity *Pr* is the probability to reject the null hypothesis *H_0_* = {*Pi_3_* = 0} against the alternative hypothesis *H_1_* = {*Pi_3_* ≠ 0}. In this case, *Pr* depends also on the pairwise interaction frequencies and we consider below two limit cases (Beagrie et al., 2017):

**Case I**:	The three loci are able to interact only in a triplet (i.e., the pairwise interaction	frequencies are all zero). In this scenario, the number of “false positives” (i.e., slices with all three loci present in a non-interacting state) is minimized.

**Case II**:	Two of the loci making up the triplet are always engaged in a pairwise interaction (i.e., pairwise probability of interaction *Pi* = 1), and the third locus can bind to the pair and form a triplet. In this case, the number of “false positives” is maximum, as it is relatively easy to capture in a single slice all three loci while they do not form a triplet (it is as easy as capturing a pair of non-interacting loci).

As the Case I and Case II give, respectively, an overestimation and an underestimation of GAM sensitivity *Pr*, we expect that the true sensitivity falls between the values calculated in these two cases (Beagrie et al., 2017).

3.2 Minimum number of tubes *m**

GAM sensitivity *Pr* depends on several technical (e.g., detection efficiency, number of nuclear profiles per tube *X_NP_*.) and biological (e.g., genomic distance, strength of interaction) factors. Importantly, *Pr* increases upon increase of sequenced tubes *m* (see **Suppl. Notes Figure SN3b**). To provide a useful guide to the experimentalist who wishes to use GAM, we calculate the minimum number of tubes *m** required to reach a sensitivity greater or equal than 90% under different combinations of GAM parameters. The optimal choice of parameters minimizes the value of *m**.

Following this approach, we estimated *m** as a function of the number of nuclear profiles *X_NP_* per tube and found that for currently used GAM parameters, *X_NP_* >1 (multiplex-GAM) typically gives higher sensitivity (**Figure 1f-h**; **Suppl. Notes Figure SN3c-d**). Moreover, our results provide a guide to choose *X_NP_* based on the values of the other parameters (eg, slice thickness) and the genomic distances (**Figure 1f,h**).

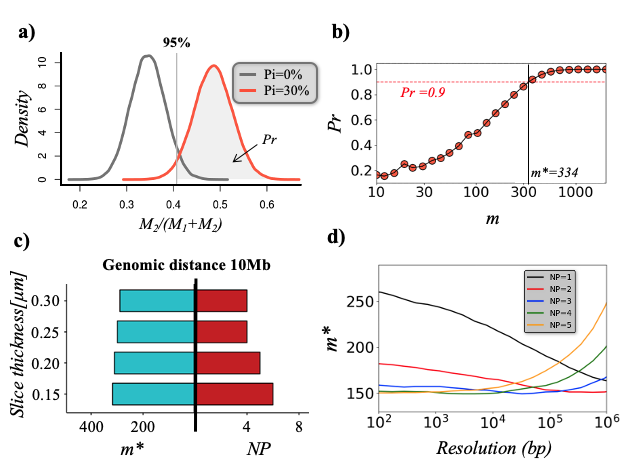

**Figure SN3**: Determining minimum number of tubes *m**.

a) The GAM sensitivity *Pr* (area under the red curve) is defined as the probability to reject the null hypothesis H_0_ = {*Pi = 0*}, and is computed as the probability of obtaining a co-segregation ratio *M_2_/(M_1_+M_2_)* greater than the 95% percentile of co-segregation ratio distribution at *Pi = 0*. The distributions, shown here as example, are computed by setting *X_NP_ = 3*, genomic distance equal to *1Mb* and detection efficiency *ε=1*.
b) By studying the sensitivity *Pr* as a function of number of tubes, we can estimate the minimum number of tubes *m** that is necessary to detect a given pairwise interaction with a sensitivity greater or equal than a threshold (represented by a dashed line), that we fixed equal to 90% for our analyses. Here, we plot, as an example, the trend of *Pr* as a function of number of tubes, by setting *X_NP_ =3*, genomic distance equal to *1Mb* and detection efficiency *ε=1*.
c) Optimal number of nuclear profiles per tube *X_NP_* and the corresponding *m** for different values of the slice thickness *h*. Similarly to **Figure 1g**, we considered pairs of loci that are at a genomic distance of 10*Mb* and interaction probability *Pi* equal to 30%*,* but we set the experimental detection efficiency ε=*83%* as estimated from the original mESC GAM dataset (408x1*NP* at *30 kb* resolution; Beagrie et al., 2017). The optimal value for *X_NP_* is consistent with the case ε=1.
d) *m** as a function of the resolution for different values of the number of nuclear profiles *X_NP_*. Here, the detection efficiency *ε* is equal to *1*, the interaction probability *Pi =* 30%, and we considered pairs at large genomic distance (≥50Mb).

For triplet interactions, a good estimation of *m** is the average of the previously described Case I and Case II, (**Section 3.1**, **Suppl. Notes Figure SN4a**). As example, the heatmap in **Suppl. Notes Figure SN4b** shows *m** as a function of the triplet interaction probability *Pi_3_* and the number of nuclear profiles *X_NP_*, for loci at 1Mb genomic distance. In this case, we find that when a small number of NPs are added per tube (i.e. 1 < *X_NP_ <* 10), few tubes are sufficient to detect frequent triplet interactions (i.e. *m** is minimized).

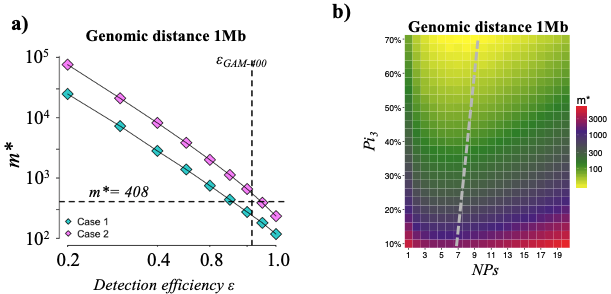

**Figure SN4:** Analysis of triplets.

a) *m** as a function of detection efficiency *ε*, where we fixed genomic distance equal to *1Mb* and interaction probability *Pi* = 50%. Here, we show the two limit cases, which give an overestimation and an underestimation of the sensitivity, respectively. The vertical dashed line marks the value of detection efficiency ε estimated from a previous GAM dataset (408x1*NP*) at 30 kb resolution, while the horizontal line marks the estimated number of tubes *m**.
b) Heatmap showing *m** as a function of the triplet interaction probability *Pi_3_* and the number of nuclear profiles *X_NP_*. We considered a detection efficiency *ε* equal to 1 and triplets at genomic distance of *1Mb*. Note that the value of *m** plotted is the average of the m* values obtained in Case 1 and Case 2. The dashed grey line indicates the trend of the minimum *m**.
4. **Analysis of Multiplex-GAM datasets**

In **Section 2.3**, we showed how SLICE model can be extended to handle Multiplex-GAM datasets, in which multiple nuclear profiles are collected in each sequencing tube (*X_NP_* >1). In this section, we test SLICE predictions on these new datasets and show that Multiplex-GAM reach a higher sensitivity. Next, we show that SLICE equations can handle combined GAM datasets with different nuclear profile per tube *X_NP_*.

4.1 **SLICE predictions on multiplex GAM datasets (*X_NP_*>1)**

In **Section 2.3**, we showed how SLICE can be extended to consider datasets with *X_NP_* >1. Through this improvement, we can estimate not only the minimal number of tubes *m** to achieve a given statistical power to reliably detect an interaction, but also the corresponding optimal value for *X_NP_,* which represents an important parameter for the new Multiplex-GAM protocol (**Section 3**). Interestingly, for a constant number of tubes *m*, we find that GAM is more sensitive when multiple nuclear profiles *NPs* are in each tube (i.e. *X_NP_*>1).

To quantitatively corroborate this prediction, we applied SLICE to two experimental datasets characterized by a different value of *X_NP_*: the new Multiplex-GAM dataset, with m=249 tubes and *X_NP_* =3 nuclear profiles per tube (briefly indicated by 249x3*NP*), and the 249x1*NP* dataset, that we generated i*n-silico* by randomly sampling a subset m=249 tubes from published data with *X_NP_* =1 (408x1*NP*; Beagrie et al., 2017) (**Suppl. Notes Figure SN5b**). First, we compute the GAM sensitivity to detect pairwise interactions as a function of their genomic distances for both datasets (**Section 3.1**). As expected, we find a higher sensitivity for the 249x3*NP* dataset (blue bars) compared to the 249x1*NP* dataset (red bars), in the considered genomic distances (**Suppl. Notes Figure SN5a**). Here, we considered a detection efficiency *ε* equal to 1 and interaction probability equal to *Pi*=30%, but these results are robust to change of these parameters. Next, we run SLICE on both datasets and estimated the prominently interacting pairs, i.e. interactions with frequencies *Pi* greater than expected value *Pi* at a chosen threshold (*P* ≤ 0.05, see Beagrie et al., 2017). By comparing the number of prominent pairwise interactions as a function of their genomic distance, we find that, as expected, more prominent interactions are detected in the 249x3*NP* dataset for all genomic distances (**Suppl. Notes Figure SN5c**), in agreement with the prediction.

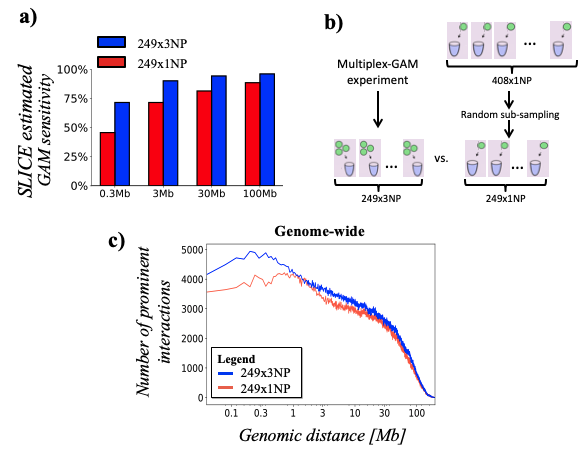

**Figure SN5.** Sensitivity of multiplex-GAM.

a) GAM sensitivity calculated by SLICE for two datasets with the same number of tubes (249) but different number of nuclear profiles per tube (*X_NP_* = 1 and *X_NP_* = 3, red and blue bars respectively), for some genomic distances. The sensitivity is predicted to be higher at all genomic distances for *X_NP_* = 3. Here we fixed a detection efficiency equal to 1 and an interaction frequency Pi = 30%, but the results are robust to changes of these parameters.
b) We compared two GAM datasets with 249 tubes from mES cells: one generated with *X_NP_*=3 (249x3NP) and another randomly sub-sampled from previously published data (Beagrie et al., 2017) with *X_NP_*=1 (249x1NP).
c) The number of prominent interactions found by SLICE is plotted as a function of the genomic distance for the 249x3*NP* (blue line) and the 249x1NP GAM data (red line). As predicted by SLICE, in the 249x3*NP* dataset a larger number of interactions are systematically identified.
4.2 SLICE predictions from combined datasets with variable *X_NP_*

We next generalized SLICE to GAM datasets including tubes with a different number of nuclear profiles. To this aim, we used a mean-field approximation, consisting of introducing a number of nuclear profiles per tube (indicated by *X_MF_*), that is the average of different number of nuclear profiles *X_NP_,* weighted with corresponding number of tubes.

As a test, we applied SLICE to the GAM-1250 dataset in which the 481x1*NP* and the 249x3*NP* datasets are combined (also indicated by 481x1*NP* + 249x3*NP*). In this case, the estimated value of *X_MF_* is:

*X_MF_* = $\frac{481x1 + 249x3}{481 + 249}$ ≈ 1.68

We run SLICE at 40 kb resolution and compared our results against the published GAM mESC-400 results at the same resolution. As expected, the detection efficiency *ε* estimated from the combined data (~86%) is comparable with the 408x1*NP* dataset at the same resolution (~93%) (Beagrie et al., 2017). Next, we compared the average probability u_0_, estimated from the experimental datasets, as a function of genomic distance (**Suppl. Notes Figure SN6a**). This is an important quantity for numerically estimating interaction frequencies *Pi* and for assessing SLICE predictions (**Section 1-2** and (Beagrie et al., 2017)). We find that the trend of u_0_ is very similar in both datasets and for all chromosomes, with Pearson correlation coefficients always around 98%.

Then, we looked at the distribution of differences of *Pi* values (**Suppl. Notes Figure SN6b**), in which, for sake of simplicity, we only show differences between values that are different from zero in both datasets. The mean value of the distribution is around zero and only low numerical differences can be found (max ~20%), proving that the two datasets have a similar distribution of *Pi*’s values. Moreover, we compared the most prominent interactions inferred from both datasets (**Section 3.1,** (Beagrie et al., 2017)). Specifically, for each chromosome we evaluated their overlap fraction (**Suppl. Notes Figure SN6c**) and computed the Pearson correlation coefficient between their common elements (**Suppl. Notes Figure SN6d**). Here, to take into account the effect of sparse matrices, we considered an element in common if the same element or at least one of its first neighbors was present.

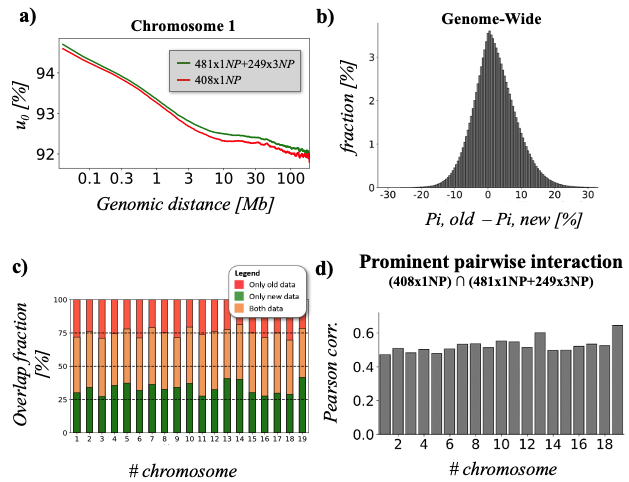

**Figure SN6**: SLICE results on Multiplex-GAM datasets.

a) The probability u_0_ of not finding neither of two loci in a not interacting state, is an important fitting parameter for SLICE (Beagrie et al., 2017). Here, we compare the estimated trend of *u_0_* as a function of their genomic distance from previous 408x1*NP* and new combined dataset (481x1*NP*+249x3*NP*) for chromosome 1. The high Spearman correlation coefficient between the two trends (about 0.97), shows that the datasets provide very similar results.
b) Distribution of differences of *Pi* values inferred at 40 kb resolution from 408x1*NP* and combined 481x1*NP*+249x3*NP* data genome-wide. For sake of simplicity, here we show differences between *Pi* values that are different from zero in both datasets.
c) Overlap of most prominent contacts between *408x1NP* and combined *481x1NP+249x3NP* datasets, at *40 kb* resolution. The overlap is around 40% between prominent *Pi*’s in the two cases.
d) Pearson correlation coefficients between the prominent *Pi* matrices computed on *408x1NP* and *481x1NP+249x3NP* datasets. Here, we correlated only most prominent contacts that are present in both datasets.
